# Generation of functional ciliated cholangiocytes from human pluripotent stem cells

**DOI:** 10.1101/2021.03.23.436530

**Authors:** Mina Ogawa, Jia-Xin Jiang, Sunny Xia, Donghe Yang, Avrilynn Ding, Onofrio Laselva, Stephanie Chin, Marcela Hernandez, Changyi Cui, Yuichiro Higuchi, Hiroshi Suemizu, Craig Dorrell, Markus Grompe, Christine E Bear, Gordon Keller, Shinichiro Ogawa

## Abstract

The derivation of mature functional cholangiocytes from human pluripotent stem cells (hPSCs) would provide a model for studying the pathogenesis of cholangiopathies and for developing novel therapies to treat them. Current differentiation protocols are not efficient and give rise to cholangiocytes that are not fully mature, limiting their therapeutic applications. Here, we describe a new strategy to generate functional hPSC-derived cholangiocytes that display many characteristics of mature bile duct cells including high levels of CFTR and the presence of primary cilia capable of sensing flow. With this level of maturation, these cholangiocytes are amenable for testing the efficacy of new cystic fibrosis drugs and for studying the role of cilia in cholangiocyte development and function. Transplantation studies showed that the mature cholangiocytes generate ductal structures in the liver of immunocompromised mice providing the first indication that it may be possible to develop cell-based therapies to restore bile duct function in patients with biliary disease.

## Introduction

The liver is comprised of many different cell types that interact to carry out the broad spectrum of functions performed by the adult organ. One of these functions, the production of bile necessary for lipid digestion is orchestrated between the hepatocytes that secrete it and the biliary system that transports it from the liver to the intestines. The biliary system consists of a series of interconnected bile ducts that are formed from a specialized population of epithelial cells known as cholangiocytes^1^. In addition to providing the structural of the bile ducts, cholangiocytes play an active role in bile flow as they are responsible for regulating its viscosity and osmolality as it moves through the liver^2^. Cholangiocytes carry out this function through a complex sensing mechanism mediated by a series of receptors, ion channels and sensory signaling molecules found on a structure known as a primary cilia that extends from the apical membrane of the cell into the lumen of the bile duct^3^. The primary cilia on the cholangiocytes function as mechano, chemo and osmo-sensors and in response to changes in bile flow or composition trigger signaling pathways that stimulate fluid secretion or absorption from the cholangiocytes^4–6^. The manipulation of bile composition through this ciliated-mediated sensing mechanism is essential for maintaining liver homeostasis^7^.

Diseases that impact cholangiocyte function are known as cholangiopathies and include monogenic disorders such as cystic fibrosis (CF) and Alagille syndrome as well as those with unknown etiology such as biliary atresia, primary biliary cirrhosis, and primary sclerosing cholangitis^8, 9^. Although these are different diseases, they share in common the fact that they all disrupt bile transport, resulting in hepatocyte toxicity, compromised organ function and in extreme cases, organ failure. The only treatment for end stage disease is organ transplantation^10^. One of the most common cholangiopathies, cystic fibrosis (CF), is caused by a mutation in the cystic fibrosis transmembrane conductance regulator (CFTR) that encodes a chloride ion channel expressed by the cholangiocyte and responsible for the regulation of bile secretion. Mutations in CFTR result in the deposition of viscous, acidic bile that is toxic to the surrounding hepatocytes, leading to liver damage and organ failure^11, 12^. As the life expectancy of CF patients increases with the availability of new treatments that improve lung function, cystic fibrosis liver disease (CFLD) is emerging as a major cause of the morbidity and mortality in these patients^11^. Small molecule drugs that rescue CFTR function have been identified and recently approved for clinical use by the FDA and other regulatory agencies^13^. These drugs have been shown to improve lung function of patients carrying the common F508del CFTR mutation, but patient-patient variability highlights an unmet need for patient-specific screening strategies for the identification of new CF drugs^14–16^. Additionally, as the drugs are largely evaluated on lung function their effect on other organs remains to be determined. Recent studies have focused on the use of CF patient organoids generated from primary intestinal, lung and nasal epithelial cells as an approach to measure both patient and tissue-specific therapeutic responses of CFTR modulators^17–19^. However, limited access to primary cholangiocytes precludes efforts to establish comparable high throughput biliary organoid screens to identify drugs that target CFTR function in these cells.

To overcome the limitation of accessibility of primary cholangiocytes, a number of groups, including ours, have turned to human pluripotent stem cells (hPSCs) as a source of these cells for modeling cholangiopathies^20–25^. In our previous study, we showed that it was possible to specify the cholangiocyte lineage from a hPSC-derived hepatoblast population through staged activation of the Notch pathway, recapitulating the observation *in vivo* that Notch signaling is required for establishing the cholangiocyte fate^21^. When cultured in semi-solid media with HGF, EGF and TGF*β*, these cells formed cholangiocyte cysts that displayed characteristics of rudimentary biliary structures including the ability to respond to agonists and drugs that activate CFTR. Other studies have similarly reported on the development of staged protocols using either monolayer or 3D culture formats to differentiate cholangiocyte-like cells from hPSCs^20, 22–25^. A number of pathways, including IL-6, EGF, TGF*β* and/Notch were manipulated at different times in the cultures, yielding cells that expressed genes and protein associated with the cholangiocyte lineage including SOX9, CK7, CK19, CFTR and SLC4A2 among others. While these studies demonstrate that it is possible to generate cholangiocyte-like cells from hPSCs, they all have limitations which include inefficient differentiation and low cell yield, difficult access to cells from semi-solid media and incomplete maturation. Notably, none of the studies to date reported the efficient development of ciliated cholangiocytes. This is a major limitation as it indicates that the cells are not mature and precludes studies on the role of cilia in normal cholangiocyte function and in the pathophysiology of different cholangiopathies.

To address this shortcoming, we developed of new monolayer-based differentiation strategy that yields high numbers of mature, ciliated cholangiocytes from different hPSC lines. Using several different screen strategies, we identify the retinoic acid, BMP, cAMP and Rho kinase pathways as key regulators of cholangiocyte maturation. The cells generated with this approach express high levels of CFTR appropriate for drug screening and contain primary cilia, that in response to fluid flow initiate intracellular Ca^2+^ release and activate CFTR. Following transplantation into the spleen of TK-NOG immunocompromised recipients, the hPSC-derived cholangiocytes migrated to the liver where they form multiple ductal structures consisting of cells that display characteristics of mature cholangiocytes. With these advances, it is now possible to engineer biliary structures with appropriate staged ciliated cholangiocytes to model different cholangiopathies and to develop new drug- and cell-based therapies to treat them.

## Results

### RA signaling promotes development of CFTR^+^ cholangiocytes

As a first step to generate mature cholangiocytes, we investigated the role of different signaling pathways on the specification of the cholangiocyte fate from day 27 hepatoblasts generated with our previously published protocol. For these studies, day 27 hepatoblasts were plated in the presence of HGF and EGF for 4 days either on OP9 jagged-1 mouse stromal cells (OP9j) to induce Notch signaling or on Matrigel without stromal support. Subsequently, agonists and antagonists of pathways known to promote early bile duct differentiation were added and the levels of CFTR, SOX9, alpha fetoprotein (AFP) and albumin (ALB) were measured following 6 days of culture. (Figure 1a and b). Cells cultured on OP9j cells without any additional factors expressed higher levels of the early cholangiocyte marker *SOX9* and lower levels of the hepatocyte markers *ALB* than those cultured on Matrigel alone, indicating that Notch signaling induced the initial stages of cholangiocyte development in the monolayer cultures. Analysis of the populations treated with the different pathway regulators revealed that only retinoic acid (RA) signaling induced significant levels of *CFTR* expression. Although expression was upregulated in cells cultured in both formats, the levels were dramatically higher (3 to 4 fold higher) in those co-cultured with OP9j. RA did not affect the levels of *SOX9*, but did further reduce the expression of *AFP* and *ALB* (Figure 1b). Western blot analyses revealed that RA concentrations of 500nM, 1μM and 2μM all induced significant levels of CFTR protein in the developing cholangiocytes (Figure 1c, d). Collectively, these findings indicate that RA signaling plays a role in the specification of the cholangiocyte lineage as demonstrated by the upregulation of *CFTR* as well as in further restricting the hepatocyte program as shown by the downregulation of the hepatoblast and hepatocyte markers *AFP* and *ALB*.

**Figure 1.**
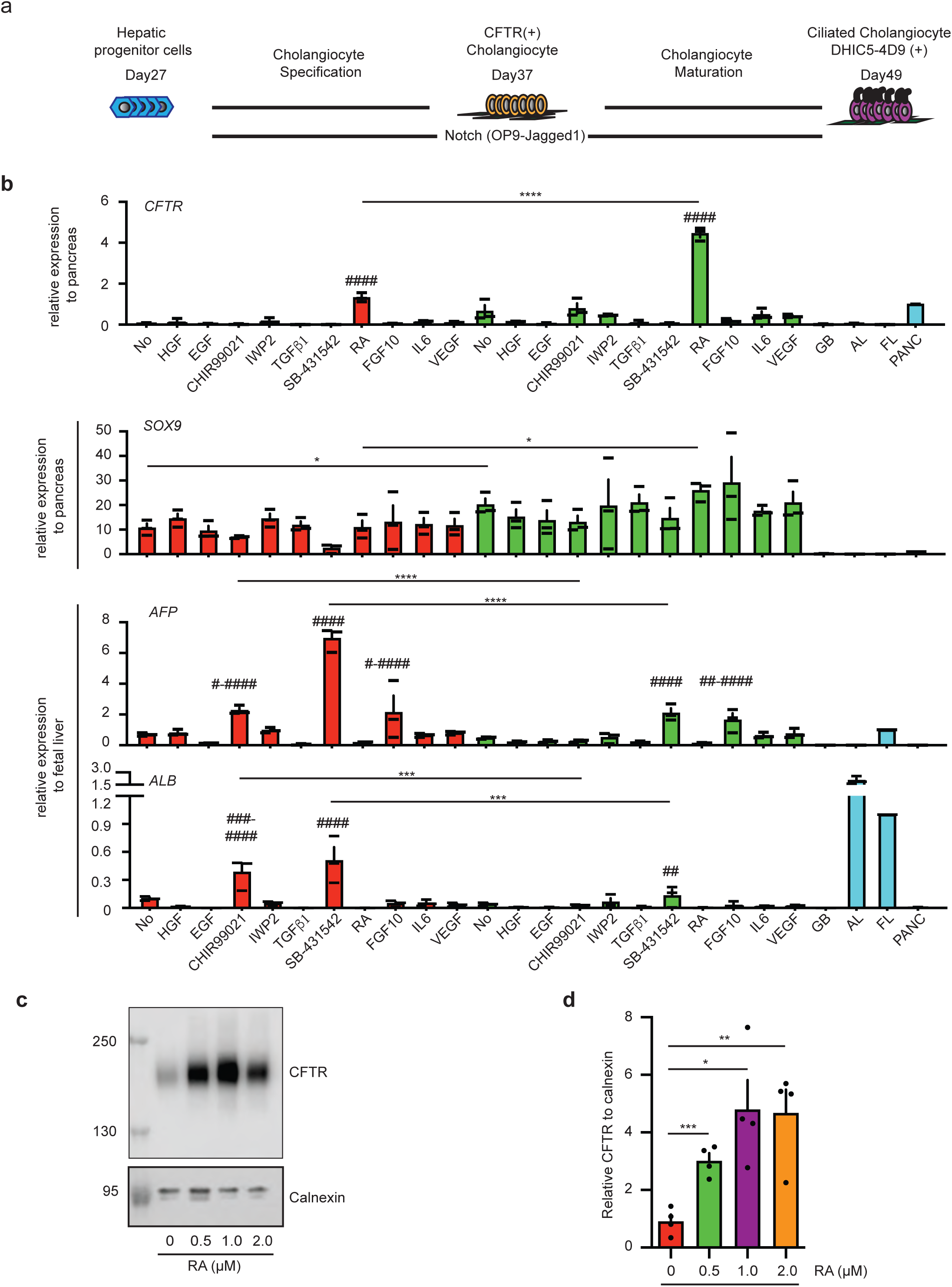
Establishment of differentiation protocol of hPSC-derived functional cholangiocytes in monolayer culture. (a) Schematic of differentiation protocol. (b) RTqPCR analysis of CFTR, SOX9, AFP and ALB expression in the presence of factors for 6 days following HGF and EGF treatment for 4 days after the passage on either Matrigel (Red) or OP9-Jagged1 (Green) from the day 27 hepatoblast. ^#^ represents statistical significance among factors in Matrigel or in OP9-Jagged1. ^#^*p* ≤ 0.05, ^##^*p* ≤ 0.01, ^###^*p* ≤ 0.001, ^####^*p* ≤ 0.0001 one-way ANOVA. * represents statistical significance between factors in Matrigel and in OP9-Jagged1 \**p* ≤ 0.05, \*\*\**p* ≤ 0.001, \*\*\*\**p* ≤ 0.0001 two-tailed Student’s t-test. Data are represented as mean ± SEM. (n=3). (c) Western blot analysis showing mature glycosylated CFTR bands in hPSC-derived cholangiocytes at day37 after treated with different concentrations of RA. (d) Quantification of mature glycosylated CFTR proteins in hPSC-derived cholangiocytes at day37 after treated with different concentrations of RA. \**p* ≤ 0.05, \*\**p* ≤ 0.01, \*\*\**p* ≤ 0.001 two-tailed Student’s t-test compared to RA 0μM. Data are represented as mean ± SEM. (n=4).

### Generation of ciliated cholangiocytes from hPSCs

Although RA promoted the development of CFTR^+^ cholangiocytes in monolayer cultures, less than 20% of the cells were ciliated, indicating that most were not fully mature. To identify regulators that would promote further maturation of the CFTR^+^ cholangiocytes, we developed a flow cytometry based screening approach using the antibody DHIC5-4D9 that stains adult human bile and pancreatic ducts^21, 26, 27^ but not the day 37 RA induced hPSC-derived cholangiocytes (Figure 2a). We analyzed agonists and antagonists of pathways known to play a role in bile duct development and maturation^28–34^ for their ability to induce the development of DHIC5-4D9^+^ cells in the hPSC-derived cholangiocyte population. As shown in Figure 2b, treatment with either a Rho-kinase inhibitor (RI), Forskolin (FSK), cAMP or the BMP inhibitor Noggin (NOG) promoted the development of DHIC5-4D9^+^ cells at day 43 of culture. The proportion of positive cells in the RI-, FSK-, cAMP- and NOG-treated groups was: 30.9 ± 4.5% 26.6 ± 4.6%, 24.3 ± 4.3%, and 18.0 ± 2.1% (mean ± SEM) respectively (Figure 2b and Supplementary Figure 1a). The combination of NOG, FSK, and RI (NFR) was more effective than each factor alone and generated populations that contained 68.0 ± 4.0% and 79.9 ± 3.7% DHIC5-4D9^+^ cells at days 43 and 49 of culture respectively (Figure 2a and Supplementary Figure 1b). The population of DHIC5-4D9^+^ cells significantly decreased in the absence of RI, suggesting that it is the main contributor to DHIC5-4D9^+^ cholangiocyte differentiation (Figure 2a). NFR did not induce the DHIC5-4D9^+^ population if RA was inhibited with BMS439, indicating that RA signaling is required for the early stages of cholangiocyte maturation and cannot be replaced by NOG, FSK, and RI (Figure 2c). As previous studies have shown that EGF signalling plays a role in the maturation of cholangiocytes from hPSCs^20–25^, we included it in our screen. EGF did not promote the development of DHIC5-4D9^+^ cells under the conditions of our cultures. Moreover, when combined with NFR, it inhibited the generation of DHIC5-4D9^+^ cells (Supplementary Figure 1b).

**Figure 2.**
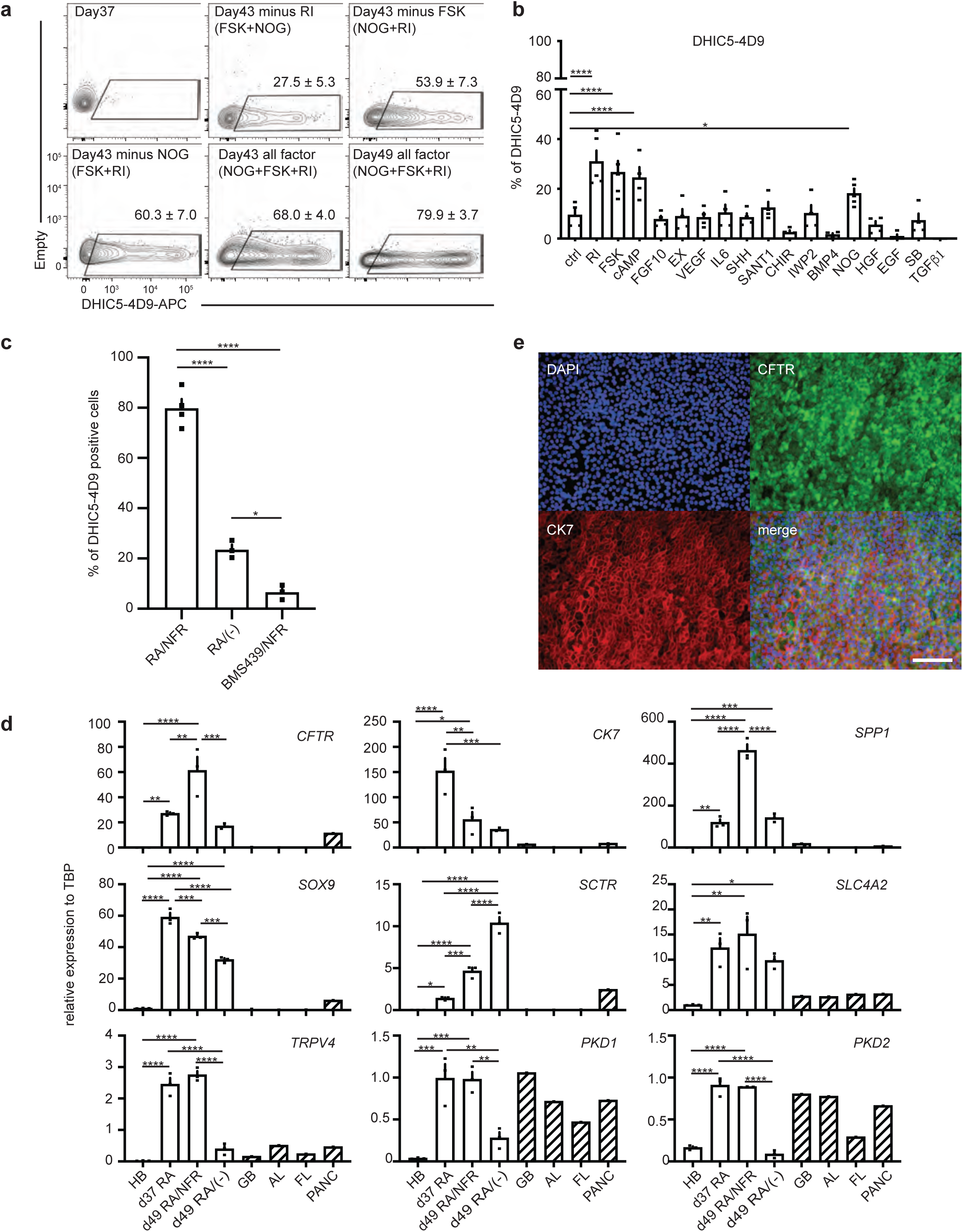
FSK, NOG, and RI promote the generation of DHIC5-4D9 positive cells. (a) Flow cytometry analysis showing the proportion of DHIC5-4D9^+^ cells at day37, day43 and day49. All factor represents addition of NFR (<u>N</u>oggin, <u>F</u>orskolin and <u>R</u>ho-Kinase inhibitor) following RA treatment. Minus represents withdrawal of each representative factor from NFR for 6 days. Data are represented as mean ± SEM (n=3-5). (b) % positivity of DHIC5-4D9^+^ cells after 6 days in the presence of factors. \**p* ≤ 0.05, \*\*\*\**p* ≤ 0.0001 two-tailed Student’s t-test compared to control. Data are represented as mean ± SEM. (n=4-5). (c) Quantification of DHIC5-4D9 positive cells in day 49 hPSC derived cholangiocytes after treatment with different indicated cytokine/small molecule combinations. \**p* ≤ 0.05, \*\*\*\**p* ≤ 0.0001 one-way ANOVA. Data are represented as mean ± SEM (n=3-4). (d) RT-qPCR analysis of the expression of the indicated gene in each population at different stages of H9 derived cholangiocytes. HB: hepatoblast, GB: gall bladder, AL: adult liver, FL: fetal liver, PANC: pancreas. \**p* ≤ 0.05, \*\**p* ≤ 0.01, \*\*\**p* ≤ 0.001, \*\*\*\**p* ≤ 0.0001 one-way ANOVA. Data are represented as mean ± SEM (n=3). (e) Immunostaining analysis showing the proportion of CFTR (green) and CK7 (red) cells in day49 cholangiocyte. Scale bar represents 200μm.

Molecular analyses revealed changes in gene expression within the RA/NFR treated population indicative of bile duct development and maturation. RA alone induced the upregulation of expression of the ductal genes *SPP1*, *CK7* and *SOX9*, of genes that encode the hormone receptor *SCTR* and the the ion channels *SLC4A2*, *CFTR,* and of those that encode proteins involved in cilia formation and function such as *TRPV4, PKD1, PKD2* in the day 37 populations (Figure 2d) . Although RA signaling promoted these changes, the levels of some of the genes, most notably those associated with cilia development and function, were not maintained over the following 12 days in the absence of NFR. The addition of NFR at day 37 led to even higher levels of *CFTR, SCTR* and *SPP1* expression by day 49 than those induced by RA alone (Figure 2d). Additionally, NFR maintained the RA-induced levels of cilia related genes in the day 49 populations. The RA/NFR induced population expressed CFTR, SOX9, CK7, ZO-1 and ASBT (bile acid transporter) proteins further supporting the interpretation that these cells represent functional cholangiocytes (Figure 2e and Supplementary Figure 2a). Flow cytometry analyses confirmed the immunostaining and molecular analyses and showed that the majority of the RA/NFR treated cells expressed CK7 and EPCAM. Very few ALB^+^ cells were detected in the population (Supplementary Figure 2b).

To demonstrate CFTR function, we measured apical chloride conductance (ACC) using a membrane potential dye (FLIPR assay) ^18^. CFTR function following the NFR treatment (day 49) was higher than that immediately following RA treatment (day 37) indicating that the cholangiocytes matured over this period of time in the presence of NFR (Supplementary Figure 2c).

To determine if the induction of the DHIC5-4D9^+^ population correlated with the development of cells with primary cilia, we stained the RA/NFR treated cells with an antibody against acetylated *α*-tubulin (Figure 3a, Supplementary Figure 2a, Supplementary Video 1). As shown in Figure 3b, the majority of cells in the population were ciliated. Quantification of different populations revealed that RA treatment alone promoted cilia development in approximately 15% of cells at day 37, and 28% of cells at day 49 in the absence of NFR. In contrast, over 85% (87.2 ± 2.5%, mean ± SEM) of the RA/NFR-treated cells were ciliated at day 49 indicating that DHIC5-4D9 staining correlated with cilia development as a parameter for cholangiocyte maturation. The addition of EGF in place of NFR did not increase the proportion of ciliated positive cells over that observed in the population treated with RA alone. Further analyses showed that the cilia on the cells expressed ARL13B, a Ras superfamily GTPase known to be present on primary cilia of mature cholangiocytes^35^ (Figure 3c and Supplementary Figure 2a). Scanning electron microscopy showed the presence of both primary cilia and microvilli, indicating that the ultrastructure of the hPSC-derived cholangiocytes was similar to of primary adult cholangiocytes^36^ (Figure 3d).

**Figure 3.**
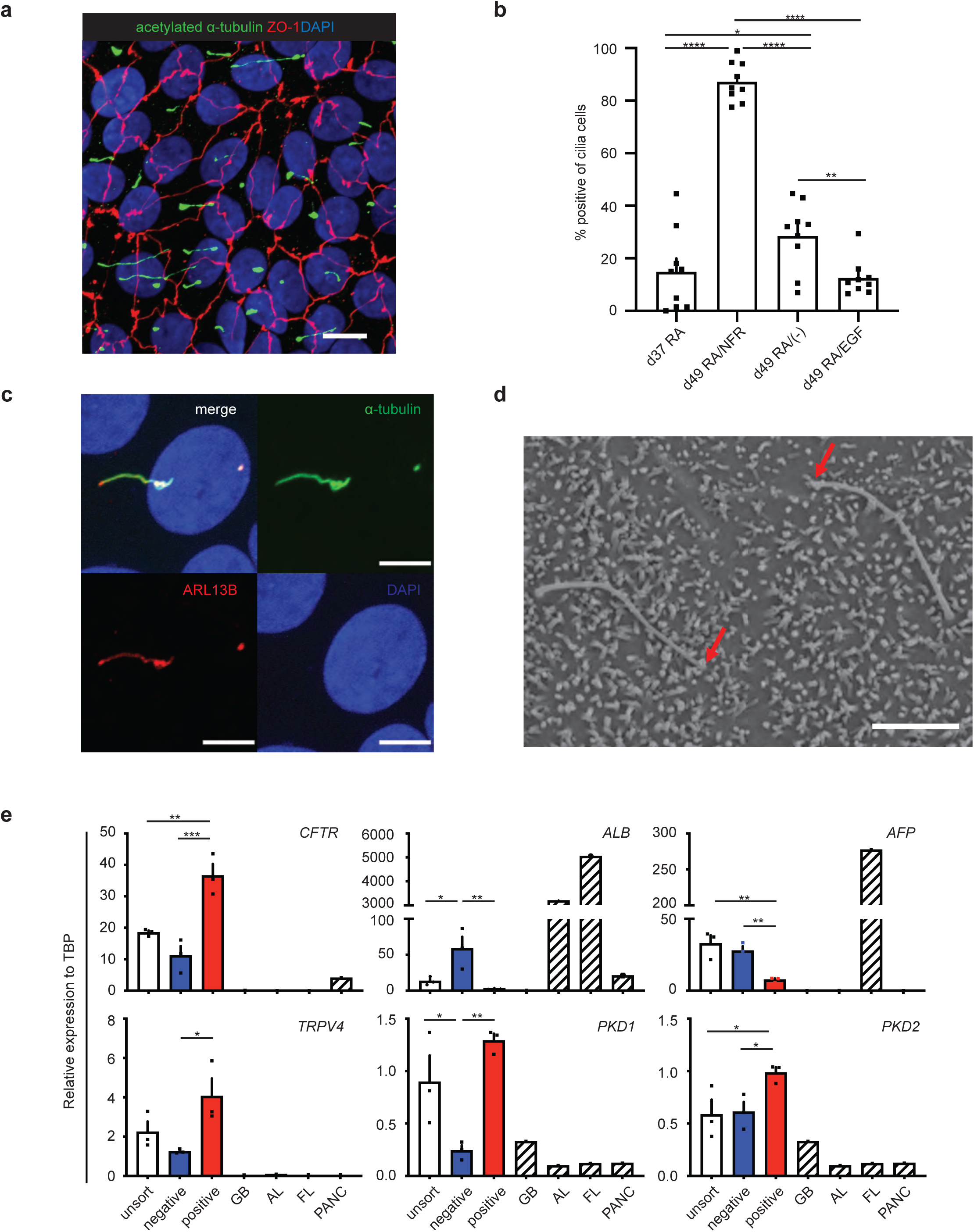
DHIC5-4D9 positive cells show primary cilia, a characteristic feature of cholangiocyte. (a) Confocal microscopy analysis showing co expression of acetylated *α*-tubulin and ZO1 in day49 cholangiocyte. Scale bar represents 10μm. (b) Quantification of cilia positive cells in day37 (after RA) and day 49 hPSC derived cholangiocytes after treatment with different indicated cytokine molecule combinations. \**p* ≤ 0.05, \*\**p* ≤ 0.01, \*\*\*\**p* ≤ 0.0001 one-way ANOVA. Data are represented as mean ± SEM (n=3). (c) High magnification image with confocal microscopy demonstrating co-localization of ARL13B with acetylated-*α*-tubulin (+) primary cilia. Scale bar represents 5μm. (d) Scanning electron microscope image of primary cilia and microvilli in 2 cholangiocytes. Red arrows indicate the tips of cilia. Scale bar represents 5μm. (e) qPCR analysis for indicated genes in flow-based sorting fractions of DHIC5-4D9^+^ cells in day49 cholangiocyte. GB: gall bladder, AL: adult liver, FL: fetal liver, PANC: pancreas. \**p* ≤ 0.05, \*\**p* ≤ 0.01, \*\*\**p* ≤ 0.001 one-way ANOVA. Data are represented as mean ± SEM (n=3).

To formally demonstrate that DHIC5-4D9 staining is a marker of maturation, we isolated the DHIC5-4D9^+^ and DHIC5-4D9^-^ fractions from a day 49 population (Supplementary Figure 3) and conducted qRT-PCR. As shown in Figure 3e, the DHIC5-4D9^+^ cells expressed higher levels of *CFTR*, *TRPV4*, *PKD1* and *PKD2* and lower levels of *ALB* and *AFP* than the DHIC5-4D9^-^ cells. These findings demonstrate that the DHIC5-4D9^+^ cells represent a more mature stage of cholangiocyte development than the DHIC5-4D9^-^ cells and in doing so validate the use of the DHIC5-4D9 antibody for monitoring cholangiocyte maturation. In addition to promoting maturation, treatment with NFR increased the yield of ciliated cholangiocytes on day 49 to an average of 2.15 ± 0.12 per day 27 hepatoblast or 11.38 ± 1.73 per input hPSC, respectively. In contrast, RA-induced populations without NFR treatment generated on average 0.05 ± 0.03 ciliated cholangiocytes per input hepatoblast on day 49 of culture (Supplementary Figure 4a). Taken together, the findings from this series of studies demonstrate that it is possible to efficiently generate functional ciliated cholangiocytes in a monolayer format with appropriate staged manipulation of specific signaling pathways.

### Measuring CF drug responses with hPSC-derived cholangiocytes

To determine if the cholangiocytes generated with the monolayer protocol can be used to evaluate the effects of modulators on CFTR function, we measured the Z prime score on H9 hPSC-derived cells using the FLIPR assay in 96 well culture formats. This analysis revealed a score of 0.478 following RA treatment (Supplementary Figure 4b), and 0.629 following NFR treatment (Figure 4a). These scores indicate that the FLIPR assay should be able to reliably predict responses to CFTR modulators designed to rescue CFTR function in cholangiocytes generated from CF iPSCs.

**Figure 4.**
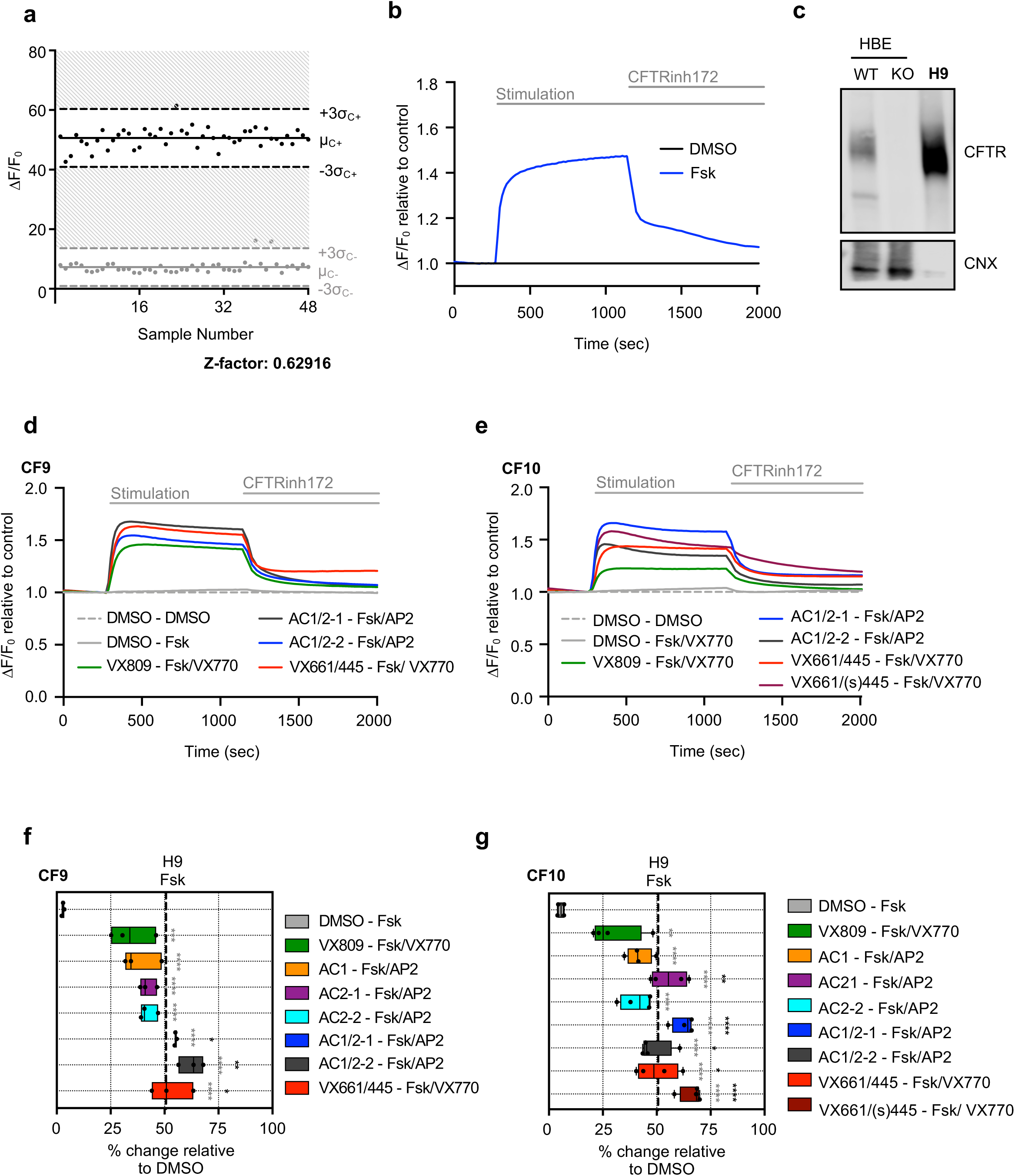
CFTR Modulator profiling of CF patient derived cholangiocytes in 96 well monolayer culture. (a) The Z factors show the quality of the FLIPR assay measuring Apical Chloride Conductance (ACC) in 96 well plate with hPSC-derived cholangiocytes at day49. (b) The kinetics of ACC by FLIPR assay in H9 derived cholangiocytes at day49. (c) Western blotting shows the mature glycosylated CFTR band in day49 H9 cholangiocyte along with the controls from CFTR expressing human bronchial epithelial HBE cell line (WT: wild type, KO: CFTR knock out). CNX: calnexin control. (d) The kinetics of ACC by FLIPR assay among different drug treatment in CF9 iPSC-derived cholangiocytes. (e) The kinetics of ACC by FLIPR assay among different drug treatment in CF10 iPSC-derived cholangiocytes. (f) Representative % value of CFTR channel activity with an exposure of DMSO and CFTR modulators in CF9 iPSC-derived cholangiocytes normalized to DMSO. Dashed line indicates FSK response in H9-derived cholangiocytes. Grey and black stars show statistical significance to DMSO and VX809/VX770 respectively, one-way ANOVA (n=3). (g) Representative % value of CFTR channel activity with an exposure of DMSO and CFTR modulators in CF10 iPSC-derived cholangiocytes normalized to DMSO. Dashed line indicates FSK response in H9-derived cholangiocytes. Grey and black stars show statistical significance to DMSO and VX809/VX770 respectively, one-way ANOVA (n=4). \**p* ≤ 0.05, \*\**p* ≤ 0.01,\*\*\**p* ≤ 0.001, \*\*\*\**p* ≤ 0.0001

To test this, we used the monolayer protocol to generate cholangiocytes from iPSCs derived from three different CF patients carrying the common F508del mutation (CF9, CF10 and GM432^37^). For comparison, the CFTR activity of cholangiocytes generated from a wild type hESC line, an iPSC line (CCRM45) and a genome edited line (CF9GC) were also evaluated. All iPSC lines had normal karyotypes and displayed characteristics of iPSCs including SSEA4, Tra-1-60, SOX2, NANOG, and OCT4 by protein expression (Supplementary Figure 5, 6, 7 and 8). These patient lines differentiated efficiently and generated populations highly enriched for CXCR4^+^cKIT^+^EPCAM^+^HDE1^+^ ^38^ endoderm by day 9 of differentiation (Supplementary Figure 9). The endoderm from each of the CF iPSC lines differentiated to give rise to hepatoblasts (Supplementary Figure 10a) and subsequently to DHIC5-4D9^+^ cholangiocytes that expressed genes indicative of mature ciliated cells (Supplementary Figure 10b and c). Staining for acetylated *α*-tubulin revealed that the RA/NFR induced populations from all of the lines contained a high frequency of ciliated cells (range 75% to 90%) indicating that the approach to promote maturation in a monolayer format is applicable to different hPSC lines (Supplementary Figure 11).

To validate the use of the *in vitro* generated cholangiocytes for drug evaluation, the patient iPSCs and control H9 hESCs were differentiated to mature cholangiocyte populations in microtiter wells, then treated with different CFTR modulators 24 hours prior to analyses of ACC using the FLIPR assay. The modulators tested include clinically used drugs (VX770, VX809+VX770, VX661/VX445+VX770) as well as emerging new CFTR modulators developed by Abbvie: AC1 (X281602), AC2-1 (X281632), AC2-2 (X300549), or combinations of them with the potentiator drug AP2 (X300529)^39^. H9 hESC-derived cholangiocytes showed a high ACC response to FSK stimulation (Figure 4b). Importantly, this response was inhibited by addition of the CFTR inhibitor CFTRinh-172, demonstrating that the assay is measuring CFTR channel activity. Western blot analysis confirmed robust expression of CFTR protein in the H9 hESC-derived cholangiocytes (Figure 4c). The cholangiocytes from the patient iPSCs showed different responses to different combinations of modulators. Although iPSCs from three different patients (CF9, CF10 and GM432) carried the same F508 del mutation, the cholangiocytes generated from them responded differently to a number of the drug combinations including VX809+VX770 that has been approved for treatment (Figure 4d, e, f, g and Supplementary Figure 12a). The triple combination of CFTR modulators AC1/AC2-1+AP2, AC1/AC2-2+AP2, or VX661/VX445+VX770 promoted better CFTR activity response than VX809+VX770 (Figure 4f, 4g, and Supplementary Figure 12a) and restored the complex glycosylated, mature form of the CFTR protein (Supplementary Figure 12b, c, d, f and g). These results demonstrate that the cholangiocytes generated with the monolayer culture described here can be used to screen for drugs that modulate CFTR function in CF patient-derived cells.

### Generation of 3D cyst/organoids from monolayer cholangiocyte populations

We previously described an approach for generating hPSC-derived 3D cholangiocyte cysts from hepatoblasts by culturing the cells in semi-solid media^21^. Although these cysts showed a Forskolin (FSK) induced CFTR mediated swelling response, the format of the cultures did not permit easy access to the cysts for further analyses. The ability to generate organoids in liquid from the monolayer cultures would represent a technically simpler and more efficient method to produce these structures. Additionally, it would also provide two complimentary platforms for measuring CFTR responses from the same population; ACC using the FLIPR assay and FSK-induced cyst swelling. To generate organoids, we dissociated day 49 cholangiocyte monolayers with collagenase to form aggregates and cultured them on a rotating platform in 10 cm non-adherent dishes. Following 6 days of culture, we observed the development of hollow cysts that were remarkably homogenous in size and shape (Figure 5a). Immunofluorescent analyses showed that cysts were comprised of CK19^+^ cells that expressed CFTR (Figure 5b).

**Figure 5.**
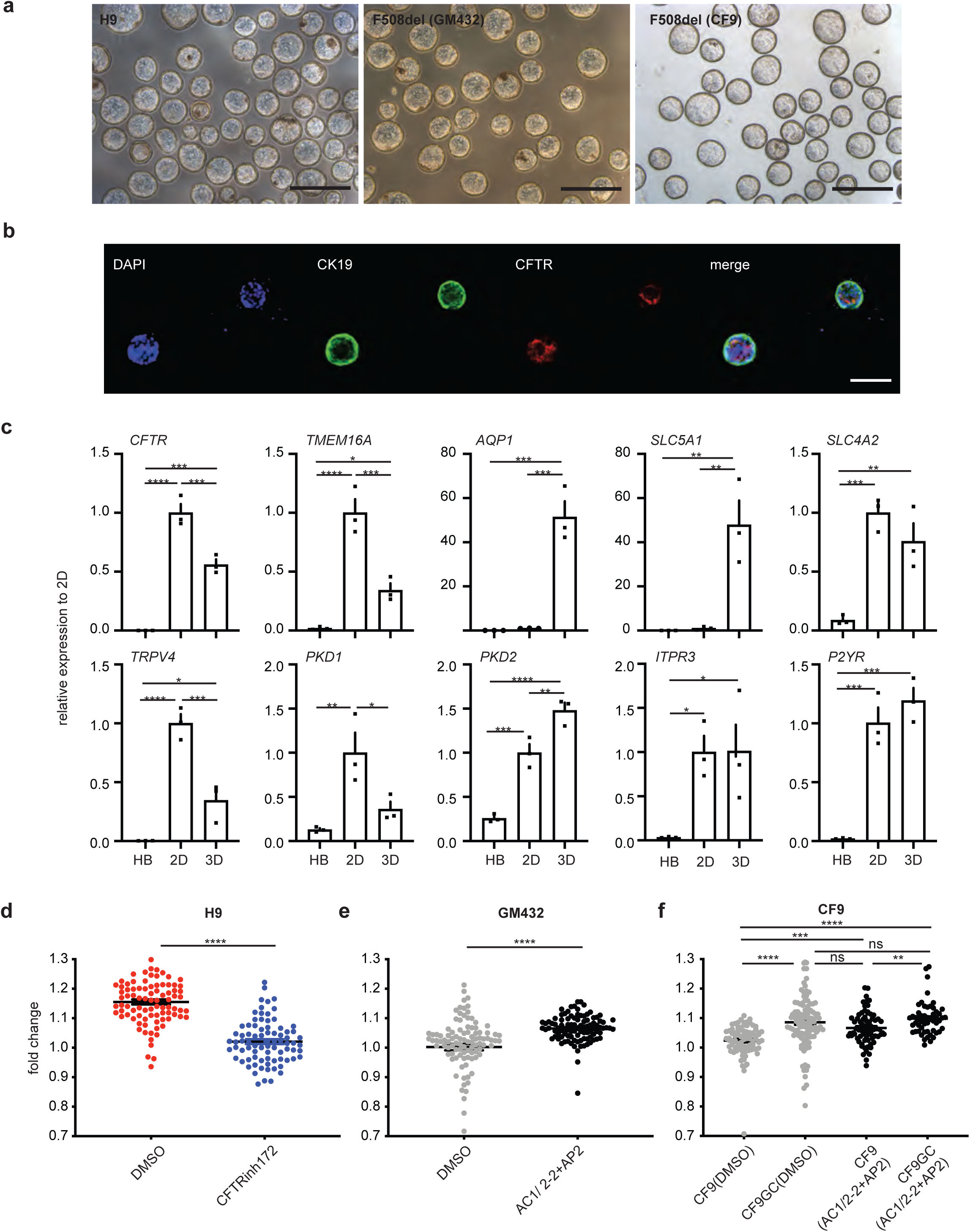
3D cholangiocytes cyst/organoids shows CFTR function and modeling of CFLD. (a) Photomicrographs of cyst/organoid structures that develop in liquid culture condition from monolayer hPSC-derived cholangiocytes. Scale bar represents 500μm. (b) Immunostaining analysis showing the co-expression of CK19 (green) and CFTR (red) in H9-derived cholangiocyte cysts. Scale bar represents 100μm. (c) RT-qPCR based expression analysis of indicated genes following 6 days culture with indicated conditions from H9-derived cholangiocyte. HB: day 27 H9 derived hepatoblasts, 2D: day55 monolayer condition, 3D cyst: day55 liquid culture condition, \**p* ≤ 0.05, \*\**p* ≤ 0.01, \*\*\**p* ≤ 0.001, \*\*\*\**p* ≤ 0.0001 one-way ANOVA. Data are represented as mean ± SEM (n=3). (d) Quantification of the degree of H9-derived 3D cholangiocyte cyst swelling 1h after FSK stimulation in the absence or presence of CFTR inhibitor CFTR Inh-172. \*\*\*\**p* ≤ 0.0001 two-tailed Student’s t-test. Data are represented as mean ± SEM (n=3). (e) Quantification of the degree of F508del (GM432)-derived 3D cholangiocyte cyst swelling 1h after FSK stimulation in the presence of DMSO or CFTR modulators. \*\*\*\**p* ≤ 0.0001 two-tailed Student’s t-test. Data are represented as mean ± SEM (n=3). (f) Quantification of the degree of F508del (CF9) or corrected F508del (CF9GC)-derived 3D cholangiocyte cyst swelling 1h after FSK stimulation in the presence of DMSO or CFTR modulators. \*\**p* ≤ 0.01, \*\*\**p* ≤ 0.001, \*\*\*\**p* ≤ 0.0001, one-way ANOVA. Data are represented as mean ± SEM (n=3).

Molecular analyses showed that the cholangiocytes in the monolayers and cysts at day 55 expressed different levels of genes that encode channel and transporter proteins essential for bile duct function. The monolayer cells expressed higher levels of *CFTR,* and the calcium activated chloride channel *TMEM16* than the cysts whereas expression of *AQP1* that encodes a water channel protein and *SLC5A1* that encodes a sodium/glucose transporter was much higher in the cysts. Expression levels of the anion exchange protein 2, *SLC4A2* and the sodium/potassium/chloride transporter *SLC12A2* (not shown) were comparable in both populations (Figure 5c).

When tested in the swelling assay, H9 hESC-derived cholangiocyte cysts showed an average 1.16 ± 0.01-fold increase in size in response to FSK within 60 minutes (Figure 5d and Supplementary Video 2). This acute response was blocked by the addition of CFTR inhibitor, confirming that the swelling was associated with CFTR function. Analyses of the GM432- and CF9 mutant-derived cysts revealed a significant swelling response (1.07± 0.01 and 1.09 ± 0.01fold respectively) to the drug combination of modulators (AC1/AC2-2+AP2), compared to the FSK treated control (Figure 5e and f). The response of the cholangiocytes from the mutation corrected CF9 cells (CF9GC) was significantly higher than that of the mutant cholangiocytes in the absence of added modulators, consistent with the presence of functional CFTR protein. The mutant cysts showed a positive swelling response to the combination of modulators. As expected, the corrected cells showed no change in response with the addition of the drug combination (AC1/AC2-2+AP2) (Figure 5f). To note, the corrected CF9 line exhibited mature glycosylated CFTR protein confirmed by western blotting (Supplementary Figure 12b). Taken together, these findings show that the cysts generated from the cholangiocyte monolayer population do show a measurable CFTR-mediated swelling response and therefore can be used together with the FLIPR assay to assess the efficacy of new CFTR modulators designed to treat CF.

### Regulation of Calcium signaling in the hPSC-derived cholangiocytes

As a further assessment of maturation, we next evaluated intracellular calcium signaling in the cholangiocytes generated from this protocol. Intracellular calcium is released in cholangiocytes in response to hormonal signalling, extracellular ATP and cilia mediated mechano-sensing^7^. The effect of ATP is important as its presence in the bile activates P2Y receptors localized at the apical membrane of cholangiocytes. Binding of ATP to P2Y receptors promotes an InsP3-dependent calcium release from intracellular stores, leading to calcium-dependent chloride efflux and bicarbonate secretion^40–43^. Q-RT-PCR analysis showed that the expression of genes related to calcium-induced fluid secretion including *P2YR1*, *ITPR3*, *TMEM16A* were upregulated, in both the day 49 monolayer cells and the 3D cholangiocyte cysts compared to hepatoblasts (Figure 5c).

To monitor calcium release in real time, we generated 3D cholangiocyte cysts from a hPSC line GCaMP3^44^, which contains a GFP fluorescent reporter targeted to calmodulin and M13 domain from myosin light chain kinase. This reporter line enables the detection of intracellular calcium release by upregulation of GFP^45–48^. Addition of ATP to the GCaMP3-derived cholangiocyte cysts induced fluorescent transients indicative of intracellular calcium release (Figure 6a, b, c, Supplementary Figure 13a, Supplementary Video 3). Aggregates of hepatoblasts which do not express P2Y1 and ITPR3 did not show this activity (Figure 6c). An ATP-induced GFP response was also measurable in the cyst-derived cholangiocytes following adherence of the cells to fibronectin coated plastic (Supplementary Figure 13b, c, d, e, and Supplementary Video 4). Under these conditions, the cysts open, enabling the cholangiocytes to attach and form a monolayer on the bottom of the chamber. These cholangiocytes also exhibited primary cilia as shown by immunostaining against acetylated *α*-tubulin (Supplementary Figure 13f). Next, we tested if flow could induce calcium release, mimicking the reaction of the primary cilia to bile flow in the liver. For these studies the 3D cholangiocyte cysts were plated in a flow chamber and media was moved over these cells using a Peristaltic Bio Mini Pump. As shown in Supplementary Figure 13g, fluid flow induced a fluorescent signal indicating that the movement of the media over the surface of the cholangiocytes resulted in the release of intracellular calcium (Supplementary Figure 13h, i, and Supplementary Video 5). Following the initial response, intracellular calcium release was observed with an additional flow stimulus (Figure 6d). The magnitude of the flow-induced response increased with flow rates (Figure 6e). We also confirmed similar intracellular calcium release in H9 and F508 del CF iPSCs-derived cholangiocytes in response to flow by Fluo-4 (Figure 6f). These findings demonstrate that the hPSC-derived cholangiocytes are able to release intracellular calcium in response to ATP and fluid flow, two pathways that regulate calcium signaling in cholangiocytes *in vivo*.

**Figure 6.**
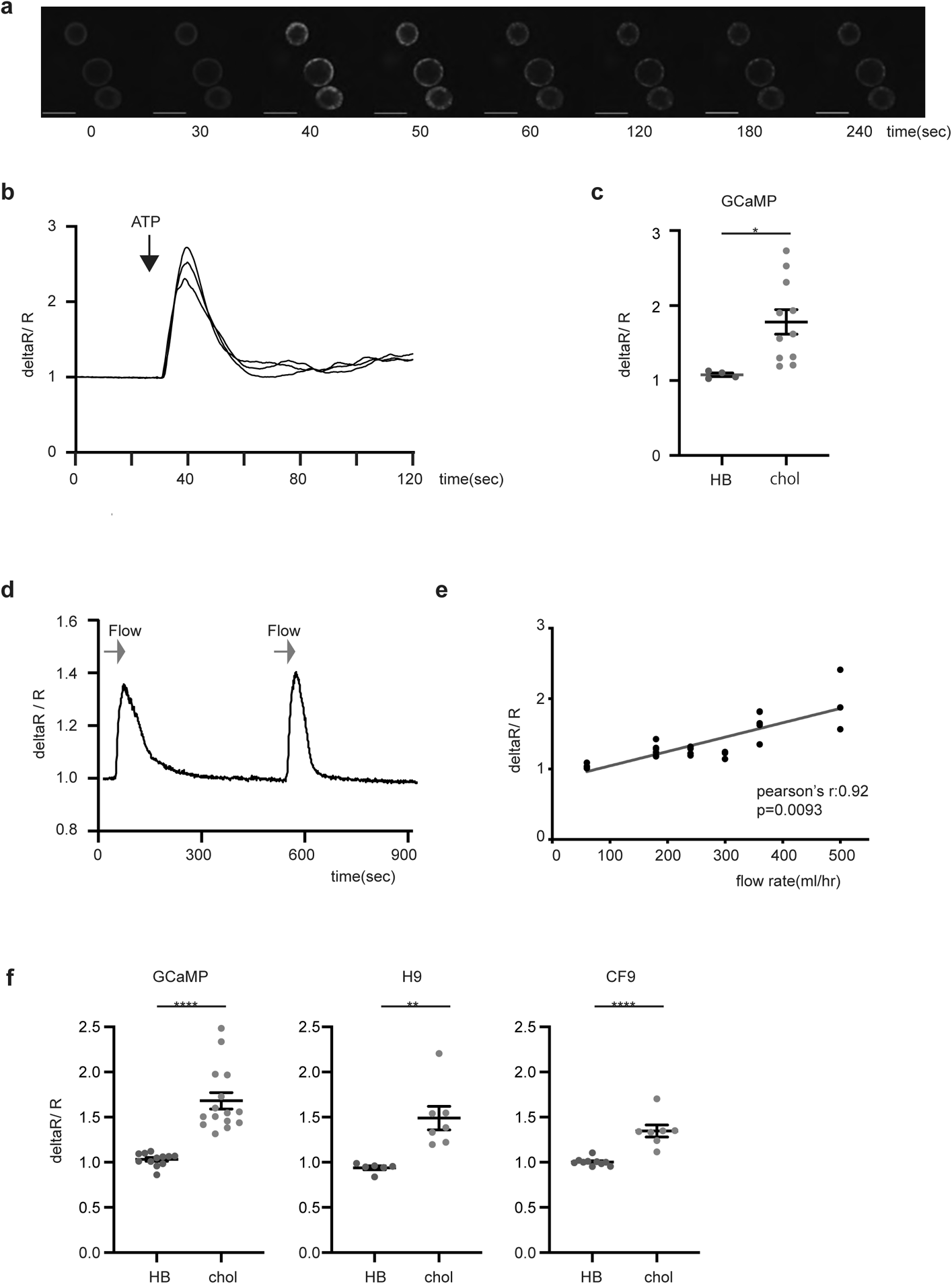
Intracellular Calcium release in ciliated hPSCs derived cholangiocyte. (a) Representative time lapse image of calcium influx in GCaMP hES derived cholangiocyte cyst (3D) in response to ATP. Scale bar represents 200μm. (b) Representative traces from Fig6 (a) showing the intra cellular calcium release in GCaMP hESC derived cholangiocytes in the response to ATP. (c) Quantification of maximum fluorescent intensity representing intra cellular Ca^2+^ release in GCaMP hESC derived hepatoblasts (HB) and cholangiocytes (chol) in the response to ATP, \**p* ≤ 0.05 two-tailed Student’s t-test (HB n=4, chol n=3). (d) Representative trace of the intra cellular calcium release in GCaMP hESC derived cholangiocytes induced by flow in the absence of ATP. (e) Quantification of maximum fluorescent intensity in GCaMP hESC derived cholangiocytes with increased flow rate (n=3). Pearson correlation coefficient was calculated by Prism8. (f) Quantification of maximum fluorescent intensity representing intra cellular Ca^2+^ release in indicated hPSC derived hepatoblasts (HB) and cholangiocytes (chol) in the response to flow, \*\**p* ≤ 0.01, \*\*\*\**p* ≤ 0.0001 two-tailed Student’s t-test (HB n=5-8, chol n=3-7).

### Flow stimulation induces CFTR functional activity via Calcium signaling in hPSC-derived cholangiocytes

Previous studies have shown that the bending of cholangiocyte cilia by luminal fluid flow in intrahepatic rat bile duct explants induced an increase in calcium and cAMP signaling, demonstrating a link between the mechanosensing properties of the cilia and intracellular signaling^3, 6^. To examine the status of mechanical stimulation of the hPSC-derived cholangiocytes, we first determined if fluid flow could induce cilia bending in these cells. As shown in Figure 7a, the cilia in the cholangiocytes did bend when exposed to flow rate of 102 μl/sec. To assess the consequence of cilia bending on the physiology of the cholangiocytes, we measured calcium signaling and ACC in GCaMP3 hPSC-derived cells subjected to fluid flow. As shown in Figure 7b, the uptake of calcium signaling and ACC activity was induced by fluid flow stimulation. Interestingly, the addition of thapsigargin, a potent inhibitor of sarco endoplasmic reticulum Ca^2+^ -ATPase that leads to a depletion of ER calcium storage, diminished fluid flow mediated increases in both calcium signaling and ACC activity (Figure 7c). These findings suggest that fluid flow mediated increases in cytosolic calcium are functionally correlated to chloride channel activity (Figure 7d). The flow-induced ACC response in H9-derived cholangiocytes were further augmented by the addition of FSK, known to activate CFTR channels through protein kinase A phosphorylation. The additive effect of flow and FSK increased the ACC response to level observed in the monolayer cultures (Figure 7e, f). Flow induced ACC was not detected in CF patient iPSC-derived cholangiocytes. However, CFTR function was rescued in the patient cholangiocytes by treatment with VX809 and significantly higher levels of rescue were observed by addition of the AC-1/AC2-2 drug combination in the presence of flow (Figure 7g, h). Taken together, these findings suggest that flow stimulation increases calcium signaling which in turn regulates CFTR channel activity and controls fluid secretion.

**Figure 7.**
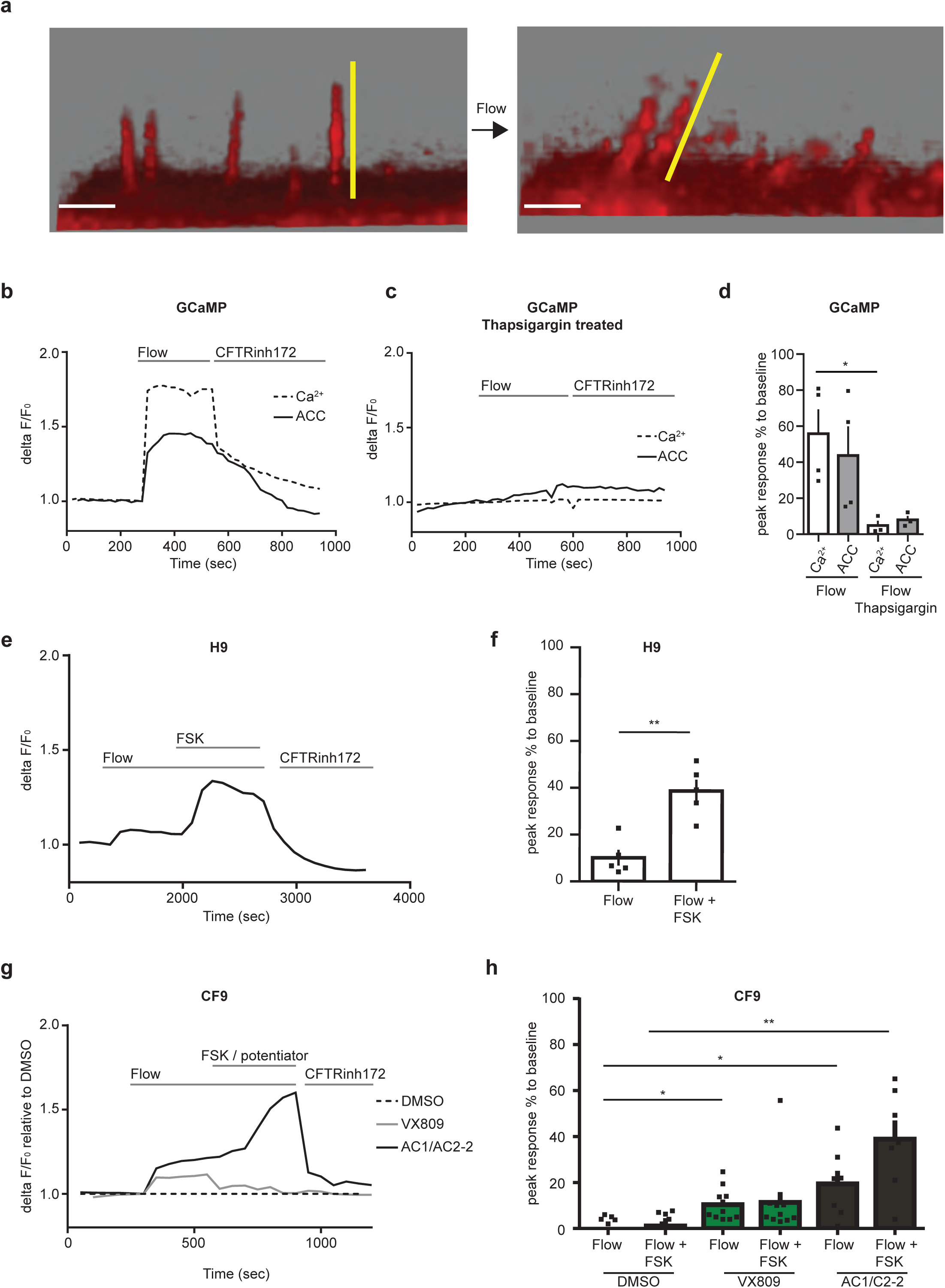
CFTR function in ciliated hPSC derived cholangiocyte in response to flow. (a) Microscopic images show that cilium extend straight from the cell surface (left). Flow (102ul/sec) bent cilium (right). Scale bars represent 5μm. (b) Correlation between calcium influx and CFTR function. Continuous flow induced calcium influx and Apical Chloride Conductance (ACC) in GCaMP derived cholangiocyte. (c) Representative trace of the Apical Chloride Conductance (ACC) and calcium influx in GCaMP derived cholangiocyte with the presence of thapsigargin in response to flow. (d) Quantitative analysis showing the peak response of calcium and ACC, \**p* ≤ 0.05 one-way ANOVA. Data are represented as mean ± SEM (n= 3-4). (e) Representative trace of the Apical Chloride Conductance (ACC) in H9 derived cholangiocyte cysts in response to flow followed by flow in the presence of FSK. Response was blocked by CFTR inhibitor172. (f) Quantitative analysis showing the peak response of ACC in H9 derived cholangiocyte cysts in response to flow, \*\**p* ≤ 0.01 two-tailed Student’s t-test (n=5). (g) Representative trace of the Apical Chloride Conductance (ACC) in CF9 derived cholangiocyte after treated with CF modulators in response to flow. (h) Quantitative analysis showing the peak response of ACC in CF9 derived cholangiocyte in response to flow in the presence of CFTR modulators \**p* ≤ 0.05, \*\**p* ≤ 0.01 one-way ANOVA. Data are represented as mean ± SEM (n= 5-11).

### Transplantation of hPSC-derived cholangiocytes

The ability to generate functional cholangiocytes from hPSCs raises the interesting possibility of developing cell-based therapies to regenerate deficient and/or diseased bile ducts in patients suffering from cholangiopathies. For this approach to be feasible, it is necessary to demonstrate that transplanted cholangiocytes can colonize the liver of a recipient and generate new ductal structures. To test this, we transplanted 10^6^ day 55 cholangiocytes into the spleen of TK-NOG mice^49^. Six weeks following transplantation, multiple duct structures consisting of cells that expressed a human mitochondria gene were detected throughout the liver (Figure 8a). The cells within the ducts also expressed CK7, CK19, and contained cilia, suggesting that they represented functional cholangiocytes (Figure 8b). As a parallel approach, we transplanted (10^6^) the mature cholangiocytes (day 55) into the subcapsular space of the kidney in NSG mice. Four weeks following the transplantation, multiple duct structures that express CK7, human mitochondria, and *α*-acetylated tubulin were also detected (Figure 8c and d). Teratomas were not observed in any of the mice that received transplants. Collectively these findings demonstrate the feasibility of generating ductal structures in the liver and ectopic sites of immunocompromised mice followed transplantation of hPSC-derived cholangiocytes.

**Figure 8.**
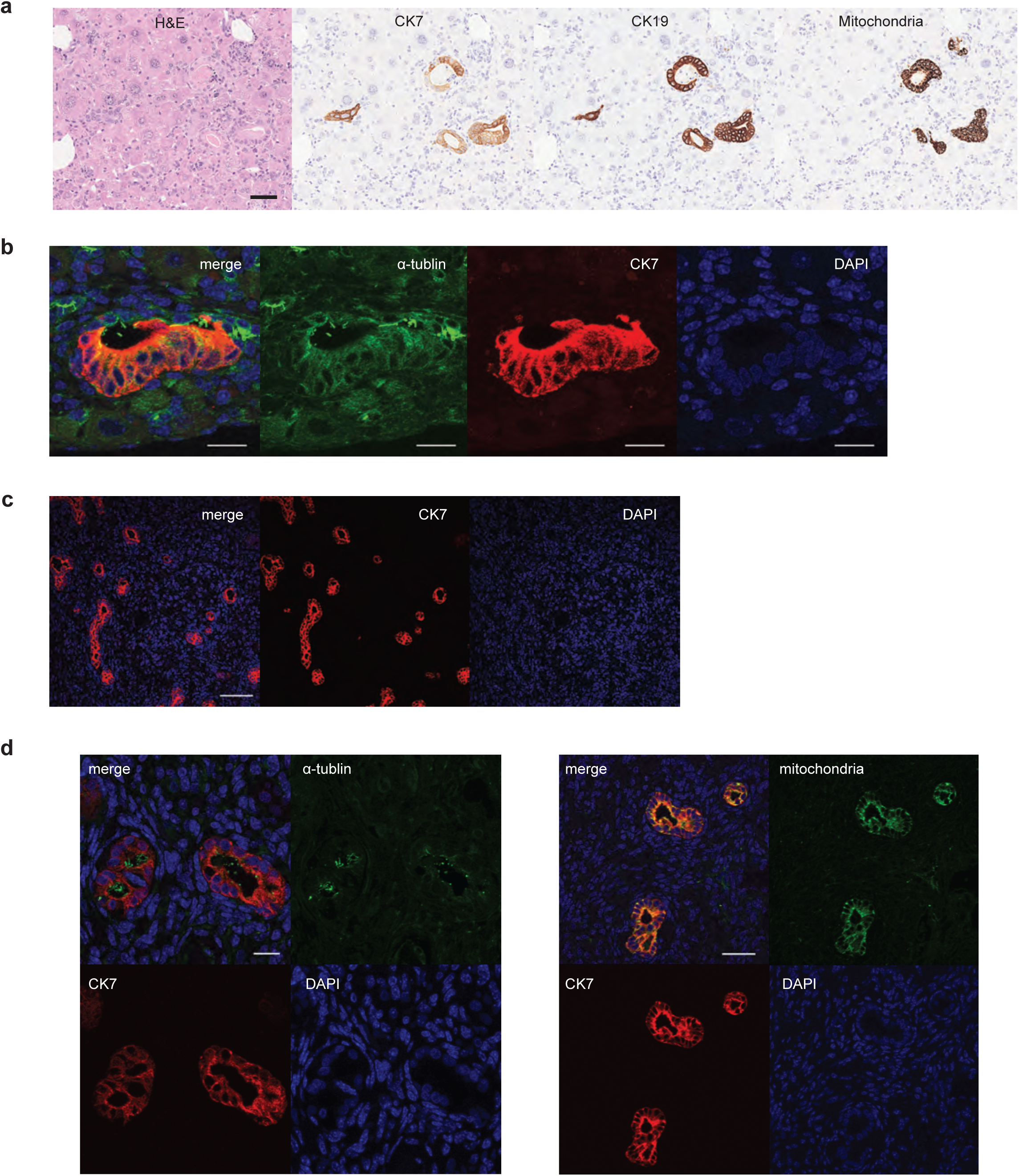
Engraftment of hPSC-derived cholangiocyte in mice liver. (a) Photomicrographs of duct-like structures generated from hPSC-derived cholangiocytes in the liver of TK-NOG mouse after 6 weeks of transplantation. HE and DAB staining with human specific antibodies. Scale bar represents 50μm. (b) Confocal image of a histological section in the liver of TK-NOG mouse after 6 weeks of transplantation showing duct structures with primary cilia (green) counterstained with CK7 (red) and DAPI (blue). Scale bar represents 20μm. (c) Confocal image of histological section showing duct structures in the kidney of NSG mouse showing human CK7 (red). Scale bar represents 100μm. (d) Left: High magnification image of confocal microscopy demonstrating that hPSC-derived cholangiocyte display co-localization of CK7 (red) with acetylated *α*-tubulin (green) in the mouse kidney subcapsular space. Scale bar represents 20 μm. Right: High magnification image of confocal microscopy demonstrating that hPSC-derived cholangiocytes display co-localization of CK7 (red) with human mitochondria (green) in the mouse kidney subcapsular space. Scale bar represents 50μm.

## Discussion

The utility of hPSC-derived cholangiocytes for therapeutic applications is dependent on the capacity of the cells to recapitulate the key functions of mature bile duct cells *in vitro*. To derive appropriate cells, it is necessary to recapitulate in the differentiation cultures the critical developmental stages and regulatory events that control their specification, expansion and functional maturation in the fetus and neonate. In our previous study, we generated hPSC-derived cholangiocytes organoid structures in semi-solid media in an attempt to recapitulate the 3D structure of the developing liver^21^. While effective, this approach has several drawbacks including inaccessibility of cells and heterogeneity of populations within the organoids. Other studies have described both 3D and 2D culture systems that promote the development of hPSC-derived cholangiocytes that display some characteristics of functional bile duct cells. None of these protocols, including the one we described give rise to functional ciliated cells^20–25^. To address this issue, we designed a monolayer based strategy that enabled the identification of signaling molecules that promote efficient maturation, yielding populations that contain a high proportion of functional ciliated cholangiocytes. These monolayer-derived cholangiocytes showed many characteristics of mature biliary cells including the presence of functional CFTR channels, the ability to mobilize calcium, responsiveness to cilia-mediated mechano-sensing and the capacity to engraft the liver of immunocompromised animals.

Previous studies in mice and the hPSC differentiation model have shown that Notch signaling plays a pivotal role in specification of the cholangiocyte fate from hepatoblasts^50, 51^. Through the use of two different screening strategies in this study, we were able to identify the retinoic acid, BMP, cAMP and Rho kinase pathways as additional regulators of human cholangiocyte development and maturation. These pathways are not redundant but rather appear to function in a stage specific pattern that establishes the framework of a regulatory roadmap of the human cholangiocyte lineage. The first step in this process is the induction of the cholangiocyte fate from hepatoblasts by Notch signaling. Patterning of the Notch-induced progenitors by RA signaling promotes the development of immature CFTR^+^ cholangiocytes that in response to manipulation of the NOG/FSK/RI pathways give rise to ciliated, functional DHIC5-4D9^+^ cholangiocytes. Although a role for these pathways has not been well defined in cholangiocyte development *in vivo*, recent studies showing that RA signaling impacts CFTR function in mouse spermatogonia stem cells^52^ supports our observations in the hPSC-derived cells. The observation that the addition of EGF reduces the proportion of ciliated cholangiocytes distinguishes our study from all others published to date that include EGF in the maturation strategy^20, 22–25^. The differences in the requirement for EGF may reflect the fact that our endpoint was ciliated cells, whereas others used different parameters including CK19, CK7, and SOX9 expression to measure cholangiocyte development.

Primary cilia are functional organelles protruding from the apical surface of many different types of epithelial cells that function to maintain tissue homeostasis by detecting changes in the extracellular environment^53, 54^. The importance of cilia to norml cell and tissue function is highlighted by the fact that mutations in genes required for ciliogenesis and primary cilia function lead to disorders/diseases known as ciliopathies^55, 56^. One such disease, polycystic kidney and liver disease is caused by mutations in either *PKD1*, *PKD2*, or *PKHD1* and is characterized by the presence of multiple cysts in the portal triad and progressive hepatic fibrosis^57, 58^. Our understanding of the role of cilia in normal cell physiology and the development of ciliopathies has been hampered by the lack of accessible model systems to study the function of these organelles. The findings presented here that flow induced calcium signaling and CFTR activation in the hPSC-derived cholangiocytes strongly suggests that the cilia are functional and capable of mechanotransduction of signals necessary for normal cell function. With these advances, it will now be possible to model ciliopathies with patient iPSCs and investigate in detail the mechanisms that lead to the onset of these diseases and identify novel therapeutics to treat them.

Over the past decade, significant progress has been made in the development of CFTR protein targeted drug therapies. While the most recent FDA approved CF drugs are able to enhance lung function in 90% of CF patients carrying the common F508del-CFTR mutation^59, 60^, there remains a number of hurdles to achieve an effective therapeutic strategy for all patients. A major issue with the drugs approved to date is that the response in different patients carrying the same mutation is variable, ranging from very modest improvement in function to an almost complete recovery of function. A second challenge is that patients with rare mutations do not respond to the drugs. Finally, the effect of the drugs is mainly measured on lung function, while the disease targets multiple organs including the pancreas, liver, and intestine. To overcome these challenges, one needs to develop assays for multiple organs from different individuals with the common F508del-CFTR mutation as well as other rare mutations. The monolayer protocol described here that gives rise to cholangiocytes with a high Z prime score provides a source of cells and culture format for measuring the effects of CF drugs on cells involved in CFLD. Indeed, we were able show that CF drugs display different degrees of rescue in these cholangiocytes generated from different patients. Additionally, with access to these mature cells, we demonstrated that it is possible to measure the effect of drugs on CFTR function in the absence of exogenous cAMP signaling, under conditions of fluid flow that were designed to mimic aspects of bile flow in the bile duct. Future studies aimed at developing organ-on chip platforms to more accurately replicate fluid flow will enable us to gain a better understanding of the interaction between cilia and CFTR function.

Although cholangiocytes represent only a small portion of the liver, they are essential for maintaining hepatocyte metabolism by controlling the viscosity through modifications of bile acid that include ion carriers and various molecules^1^. Diseases that affect cholangiocyte function such as primary sclerosing cholangitis, primary biliary cirrhosis, CFLD, and biliary atresia can impact the viscosity and flow of bile acid, resulting hepatocyte toxicity^61^. Regardless of the etiology of disease, the only effective current treatment for advanced stages of these cholangiopathies is liver transplantation. As the shortage of donor organs limits the number of patients that can be transplanted, other interventions such as cell replacement therapy need to be developed to treat patients with cholangiopathies. A recent study in mice has shown that it is possible to regenerate biliary structures from transplanted cells in the mouse model of bile duct paucity. In this case, adult hepatocytes were shown to have the potential to generate new cholangiocytes and connect to the existing biliary network in the mouse liver^62^. Our findings showing that following transplantation, hPSC-derived cholangiocytes can migrate from the spleen to the liver and form ductal structures adds additional support for the concept of cell-based therapy to treat cholangiopathies. Current studies are aimed at determining if these cholangiocyte derived ductal structures are functional.

In summary, we report on the successful generation of functional ciliated cholangiocytes from hPSCs that will provide new opportunities to study and treat diseases of the liver. Both monolayer- and organoid-derived cells are adaptable for further bioengineering approaches to produce more complex hPSC-derived liver tissue with biliary ductal structures. Additionally, the successful engraftment of differentiated cholangiocytes in the liver of immune-incompetent mice provides the basis for developing novel therapeutic strategies to treat cholangiopathies.

## Methods

### Human ES and iPS cells differentiation to cholangiocytes in monolayer

Human ES/iPS cells were differentiated to hepatoblasts as described previously^21^ with slight modification (manuscript is prepared). 30 Gray irradiated OP9 jagged-1 (OP9j) cells were plated on 2.5% Matrigel coated wells (12 well plates) at a concentration of 200,000 cells per well of 12 well plate in alpha-modified minimum essential media (a-MEM) supplemented with glutamine (2mM) and 20% fetal bovine serum. To induce cholangiocytes differentiation, day 27 hepatoblasts were dissociated by TrypLE and then plated onto the irradiated OP9j cells. The plated cells were cultured in DMEM/Ham’s F12 (1:1) (DMEM/F12) media supplemented with 0.1% BSA, 1% vol/vol B27 supplement, ascorbic acid, Glutamine, MTG, HGF (20 ng/ml), and Epidermal growth factor (EGF: 50 ng/ml) for 4 days. To induced CFTR expression in cholangiocyte-like cells, following HGF and EGF treatment, the medium was switched into DMEM/F12 medium with 0.1% BSA, 1% vol/vol B27 supplement, ascorbic acid, Glutamine, MTG, and Retinoic Acid (1μM: Sigma) for another 6 days. To promote the maturation of cholangiocytes that express primary cilia and DHIC5-4D9, the cells were cultured with DMEM/F12 medium with 0.1% BSA, 1% vol/vol B27 supplement, ascorbic acid, Glutamine, MTG, Noggin (50ng/ml), ROCK inhibitor Y-27632 (5μM) and Forskolin (FSK:5μM) for 12 days. For iPSCs, day 27 to 33 hepatoblasts were differentiated to cholangiocytes due to the slow kinetics measured by flow cytometric ALB positivity. More than 85% of ALB positive hepatoblasts were used for the further cholangiocyte differentiation. The medium for all steps of cholangiocytes differentiation were changed every two days. The differentiation was maintained in an ambient O2 incubator.

### Generation of 3D cholangiocyte organoids

Day 49 cholangiocyte following the differentiation in monolayer were dissociated with collagenase type I and then small clumps of cholangiocyte cells were plated on low attachment cluster dishes and cultured with the same medium for monolayer differentiation. The 3D cholangiocyte organoids spontaneously formed cyst-like structures within 6 days. The differentiation was maintained in an ambient O2 incubator.

### WT iPSC, CF patient iPSCs and mutation corrected iPSCs

4 iPSC lines were obtained from the CFIT Program (https://lab.research.sickkids.ca/CFIT/63). CCRM45 is WT iPSC. Two CF lines (CF9 and CF10) are carrying the common CF mutation F508del. CF9GC is the mutation corrected iPSC, generated with the CRISPR-Cas9 technology from the CF9. CCRM45, CF9, CF10, and CF9GC were generated at the Centre for Commercialization of Regeneration Medicine (CCRM) in Toronto.

### Flow cytometry

Differentiated cells were dissociated into single-cell suspensions. Dead cells were excluded during flow cytometry analyses and gating was determined using isotype control. For cell surface marker analyses, staining was carried out in PBS with 10% FCS. For detection of intracellular proteins, staining was performed on cells fixed with 4% paraformaldehyde (PFA: Electron Microscopy Science, Hatfield, PA, USA) in PBS. Cells were permeabilized with 90% ice-cold methanol for 20 min for ALB, AFP, and CK7 as previously described^21^. Cells were subsequently incubated with secondary antibodies for 30 min at room temperature. The stained cells were analyzed using LSR Fortessa flow cytometer (BD). FACS antibodies and their dilution ratio are listed in Supplementary Table 1 and 2.

### Immunostaining

For staining of monolayers, cells were fixed with 4% PFA for 15 min and permeabilized with 0.2% Triton X-100 or cold 100% methanol (CFTR, ASBT) before blocking. Cells were washed three times with PBS for 10 min at room temperature (RT) before and after each staining step. All antibodies for monolayer were diluted in DPBS + 0.1%BSA + 0.1% TritonX-100. For the staining of the 3D cyst structures, samples were fixed with 4% PFA and washed with DPBS and permeabilized with 0.2% Triton X-100 or cold 100% methanol (CFTR) before blocking. Cells were washed three times with PBS for 10 min at RT before and after each staining step. Antibodies for 3D cysts were diluted in DPBS + 0.3%BSA + 0.3% TritonX-100. DAPI was used to counterstain the nuclei. Monolayer and 3D cysts staining were visualized using a fluorescent microscope (Leica CTR 6000) and images were captured using Leica Application software. For the detection of primary cilia/ ZO-1 in monolayer cultures, the differentiation was carried out on matrigel -coated coverslips, and images were visualized using the NIKON A1 Resonant Confocal Microscope and images were captured using the Nikon Elements software. Paraffin-embedded sections were dewaxed with xylene, rehydrated, placed in Tris-EGTA-buffer (TES: 10mM Tris, 0.5mM EGTA, pH9.0) and subjected to heat-induced epitope retrieval for 20 min before blocking. All antibodies were diluted in DPBS + 0.3%BSA + 0.3% TritonX-100. Paraffin-embedded sections were analyzed using a confocal fluorescence microscope (NIKON A1 Resonant Confocal Microscope) and images were captured using the Nikon Elements software. Antibodies are listed in Supplementary Table 1 and 2.

### Scanning electron microscopy (SEM)

After standard SEM tissue preparation, the surface morphologies of the hPSC-derived cholangiocytes were sputter coated with gold-palladium prior to loading in the SEM. Images were observed using a SU3500 scanning electron microscope (Hitachi, Japan) with a 5kV magnification.

### Quantitative real-time PCR

Total RNA was prepared using RNAqueous Micro Kit (Invitrogen) and treated with RNase-free (Ambion). 500ng to 1ug RNA was reverse-transcribed into cDNA using iSCRIPT Reverse Transcription Supermix (BIO RAD). qPCR was performed on a C1000 Touch Thermal Cycler (BIO RAD) using a SsoAdvanced Universal SYBR Green Supermix (BIO RAD). Expression levels were normalized to the housekeeping gene TATA box binding protein (TBP) as previously described^21^. Oligonucleotide sequences are available in Supplementary Table 3. Control samples of RNA of adult liver (AL), fetal liver (FL), gall bladder (GB), and pancreas (PANC) are listed in Supplementary Table 4.

### Apical chloride conductance (ACC) assay

Apical CFTR-mediated chloride conductance was measured as previously described^18^. Briefly, hPSC-derived cholangiocytes were grown in 96-well plates and treated for 24h with either or combination of CFTR modulators, 3μM VX809, 3μM VX661 (Selleck), 3μM R and S-VX445 (MedChemExpress), 0.5μM AC1 (X281602), 3μM AC2-1 (X281632), 3μM AC2-2 (X300549) (Abbvie), or DMSO. Cells were labelled using blue membrane potential sensitive FLIPR dye dissolved in sodium and chloride-free buffer (150mM NMDG, 150mM gluconolactone, 10mM HEPES, pH 7.4, 300mOsm) at a concentration of 0.5mg/ml. Cells were incubated for 30 min at 37°C. The plate was read in a fluorescence plate reader (FLIPR® Tetra System or SpectraMax i3; Molecular Devices) at 37°C. After reading baseline fluorescence, CFTR was stimulated with a combination of the cAMP agonist Forskolin (10μM) and potentiators, 1μM VX770 (Selleck) or 1.5μM AP2 (X300529) (Abbvie). CFTR-mediated depolarization was detected as an increase in fluorescence, and repolarization was detected as a decrease with addition of 10μM CFTR-specific inhibitor CFTRInh-172 to all wells.

### Western Blotting

Cells were lysed in modified radioimmunoprecipitation assay buffer (50 mM Tris-HCl, 150 mM NaCl, 1 mM EDTA, pH 7.4, 0.1% SDS, and 0.1% Triton X-100) containing a protease inhibitor cocktail (Roche) for 10 min, and the soluble fractions were analysed by sodium dodecyl sulfate-polyacrylamide gel electrophoresis (SDS-PAGE) on 6% Tris-Glycine gels (Life Technologies, Carlsbad, CA)^64^. After electrophoresis, proteins were transferred to nitrocellulose membranes and incubated in 5% milk. CFTR bands were detected with human CFTR IgG2b mAb596 (1:2000, University of North Caroline, Chapel Hill, NC, code: A4, Cystic Fibrosis Foundation Therapeutics Inc.) overnight at 4°C and horseradish peroxidase-conjugated goat anti-mouse IgG secondary antibody (1:5000, Pierce, Rockford, IL) for 1 h at RT. Calnexin bands were detected with calnexin-specific rabbit pAb (1:5000, Sigma, St. Louis, MO, cat. no.: C4731) primary antibody overnight at 4°C and horseradish peroxidase–conjugated goat anti-rabbit IgG secondary antibody (1:5000, Pierce) for 1 h at RT. Wt CFTR and calnexin bands were exposed using Amersham ECL Western Blotting Detection Reagent (GE Healthcare Life Sciences, Canada) for 1-5 min exposures on the Li-Cor Odyssey Fc (LI-COR Biosciences, Lincoln, NE). Relative expression level of CFTR proteins were quantified by densitometry of immunoblots, using Image Studio Lite software (Version 5.2.5) from Li-Cor Odyssey Fc (LI-COR Biosciences)^65^.

### CFTR swelling assay

The CFTR swelling assay was performed as described previously^21^. Day 6 to day 12 3D cholangiocytes were replaced in 1 well of 384 well plate with Hank’s buffered solution and imaged using live cell imaging at 37 °C (ZEISS Axio Observer). cAMP agonists-Forskolin (10μM) were added and images were taken every 15 min for 1h. One well in each experiment was pre-incubated with 50μM CFTR-specific inhibitor-Inh172 (Selleck Chemicals) or DMSO control for 1h. CF-patients and mutation corrected CF iPSC-derived cholangiocytes were treated with either 3μM VX809 (Selleck), or 0.5μM AC1 and 3μM AC2-2 (Abbvie) for 24 hrs before the assay. CFTR was stimulated with the cAMP agonist FSK (10μM) and potentiators, 1μM VX770 (Selleck) or 1μM AP2 (Abbvie). The total area of each cysts was calculated before and 1h after stimulation using ImageJ (NIH-version 1.51). The area of each cyst after 1h stimulation was calculated and plotted. 20-40 cysts structures from each well were measured for each experiment (n=3).

### Calcium signaling assay

GCaMP 3 (gift from Michael Laflamme Lab, Toronto) and H9, CF9 iPSC were differentiated to cholangiocyte cysts and plated down in Lab-Tek Chamber Slide (Millipore) or in flow chamber (ibidi) 2 days prior to the assay. Day 27 hepatoblasts were aggregated and plated down 2 days prior to the assay. Chambers were pre-coated with fibronectin. The cells were replaced in Tyrode buffer before the assay. H9 derived-cholangiocytes and CF9 derived cholangiocytes were stained with 10μM Fluo4 (Invitrogen) with 0.04% Pluronic F127 (Invitrogen) for 30 min prior to the assay. To measure calcium response to ATP in 3D cysts, GCaMP derived cholangiocyte 3D cysts were embedded in the type1 collagen gel just before the assay. Calcium response to 20 μM ATP (Sigma) or flow (Perista BioMini Pump: ATTO) were analyzed using a confocal fluorescence microscope (NIKON A1 Resonant Confocal Microscope) and images captured using the Nikon Elements software.

### ACC assay under the flow

Cholangiocytes were prepared and plated down the same way as described in calcium signaling assay. CF patient derived cholangiocyte were treated as the same concentration of CFTR modulators and stained with FLIPR solution as mentioned in ACC assay. 2 μM Thapsigargin (Sigma) was used for calcium inhibition. Images were acquired using Leica SP8/STED confocal microscope. Data were processed with Volocity software suite (PerkinElmer).

### Mice and transplantation

TK-NOG mice transplanted at the Central Institute for Experimental Animals (CIEA) in Japan. Ganciclovir (10mg/kg) was administered into 7-8 weeks old adult TK-NOG mice 7 to 10 days prior to the transplantation. The details of induction of liver injury and transplantation were mentioned previously^49, 66^. NSG mice purchased from The Jackson Laboratory (Bar Harbor, ME) were kept in a specific pathogen-free mouse facility at the Princess Margaret Cancer Research Tower (PMCRT) and were used at 8-10 weeks of age. Experiments were performed in accordance with the protocol approved by University Health Network Animal Care Committee. Day 55 monolayer cholangiocytes were dissociated by trypsin-EDTA and one million cells were injected into the spleen of TK-NOG mice (CIEA) or the kidney subcapsular of NSG mice (PMCRT). At the indicated time points following after the transplantation, animals were euthanized, and their tissue was isolated and fixed with 10% formalin. Fixed samples were embedded in paraffin, sectioned, and stained with hematoxylin and eosin (H and E) for morphological analyses. Immunohistochemistry and DAB staining were performed as described previously^21^.

### Statistics

Statistical methods and the numbers of biological replicates were indicated in figure legends. SEM was calculated using data from biological and technical replicates.. p<0.05 was considered statistically significant (* or # <0.05, ** or ## <0.01, *** or ### <0.001, **** or #### < 0.0001). Statistical analysis was performed using GraphPad Prism 8.

## Supporting information

Supplemental figure Ogawa

## Acknowledgements

We thank Dr. Michael Laflamme for providing GCaMP cell line, and Abbvie for providing the reference tool compounds AC1, AC2-1, AC2-2 and AP2. iPSC lines were obtained through the CFIT Program (https://lab.research.sickkids.ca/CFIT), which is funded by the SickKids Foundation and Cystic Fibrosis Canada. We thank CCRM (Centre for Commercialization of Regenerative Medicine) especially Dr. Linse Munsie for the technical service to establish iPSC lines. This work is supported by the CFIT Program (CEB), and JSPS KAKENHI grant number JP18K08589 (SO). This research is part of the Government of Canada through Genome Canada and the Ontario Genomics Institute (OGI-148) (awarded to CEB, GK and SO), and part of the University of Toronto’s Medicine by Design initiative, funding from the Canada First Research Excellence Fund (awarded to CEB, GK and SO).

## Author contributions

MO and SO performed experimental design, data acquisition, analysis, interpretation and wrote the paper. JJ, SX, and DY performed experiments and data analysis. AD, MH, CC, OL, SC, and YH performed experiments. CD provided technical support. HS, and MG provided experimental guidance. CEB, GK, and SO supervised experiments, edited, revised and performed the final approval of the paper.

