## Supplemental figure Ogawa for "Generation of functional ciliated cholangiocytes from human pluripotent stem cells"

Ogawa et al.

a

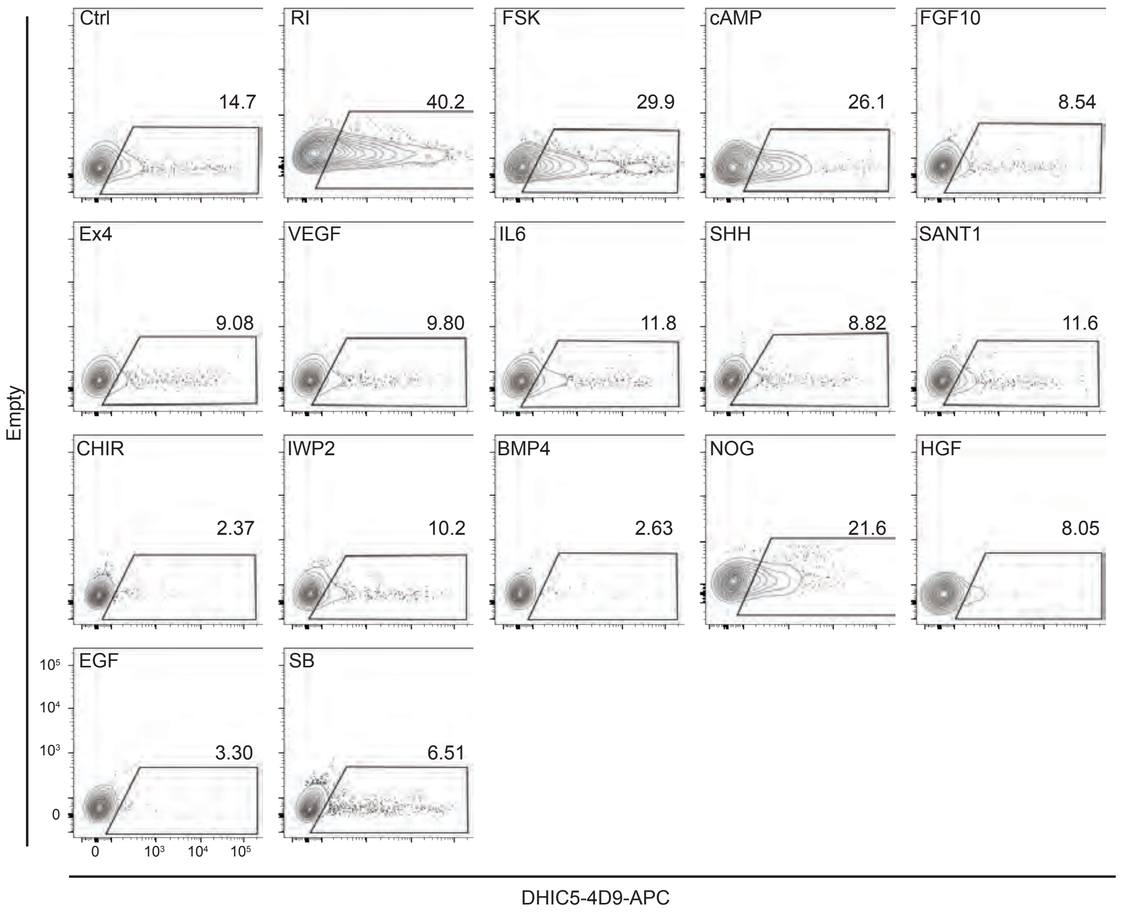

b

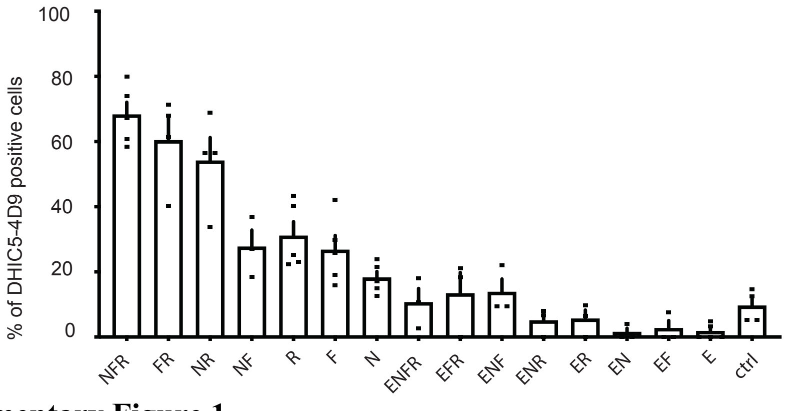

**Supplementary Figure 1**

**Identification of factors which upregulate DHIC5-4D9 expression by Flow cytometry analysis of hPSC-derived cholangiocytes.**

(a) Representative flow cytometry analysis screening of the upregulation of DHIC5-4D9 expression in day 43 hPSC derived cholangiocytes after treatment with different indicated cytokines.

(b) Quantification of DHIC5-4D9 positive cells in day 43 hPSC-derived cholangiocytes after treatment with different indicated cytokine combinations (n=3-5). Data are represented as mean ± SEM.

a

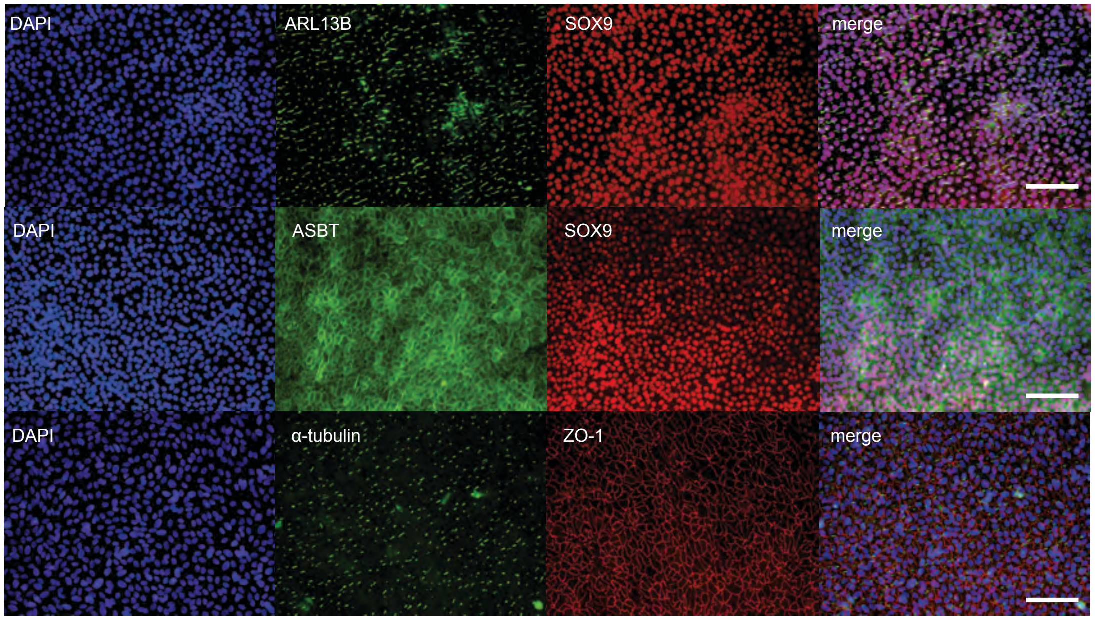
DAPI
ARL13B
SOX9
merge
DAPI
ASBT
SOX9
merge
DAPI
α-tubulin
ZO-1
merge

b

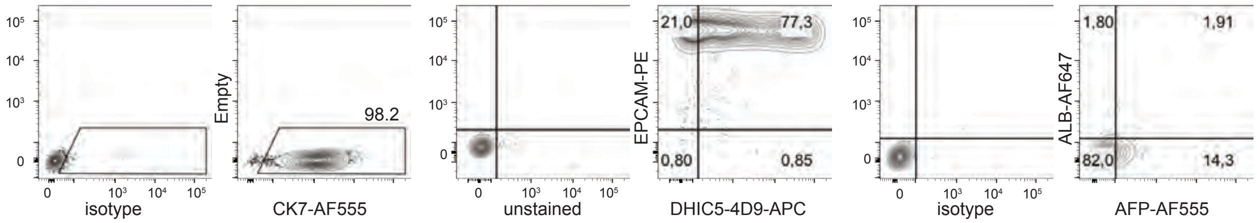
isotype
Empty
98.2
CK7-AF555
unstained
EPCAM-PE
21,0
77,3
0,80
0,85
DHIC5-4D9-APC
isotype
ALB-AF647
1,80
1,91
82,0
14,3
AFP-AF555

c

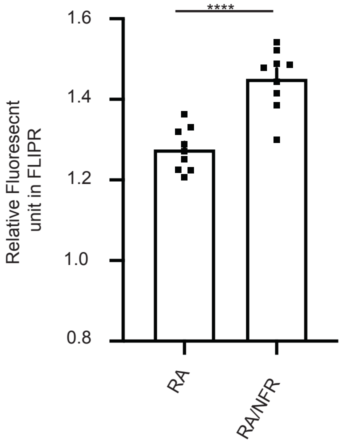
Relative Fluoresecnt unit in FLIPR
****
1.6
1.4
1.2
1.0
0.8
RA
RA/NFR

**Supplementary Figure 2**
**Characterization of hPSC-derived cholangiocytes.**

(a) Immunostaining images of day49 cholangiocytes demonstrating ARL13B (green) and SOX9 (red) in upper panel, ASBT (green) and SOX9 (red) in the middle panel, acetylated $\alpha$-tubulin (green) and ZO-1 (red) in the bottom. Scale bar represents 200μm.

(b) Representative flow cytometry analysis showing CK7 (left), EPCAM and DHIC5-4D9 (middle), and ALB and AFP (right) positivity in day49 hPSC-derived cholangiocytes.

(c) Quantification of the maximum intensity of Apical Chloride Conductance (ACC) in day 37 and day49 cholangiocytes (n=4). Data are represented as mean ± SEM. $^{****}p \leq 0.0001$ two-tailed Student's t-test.

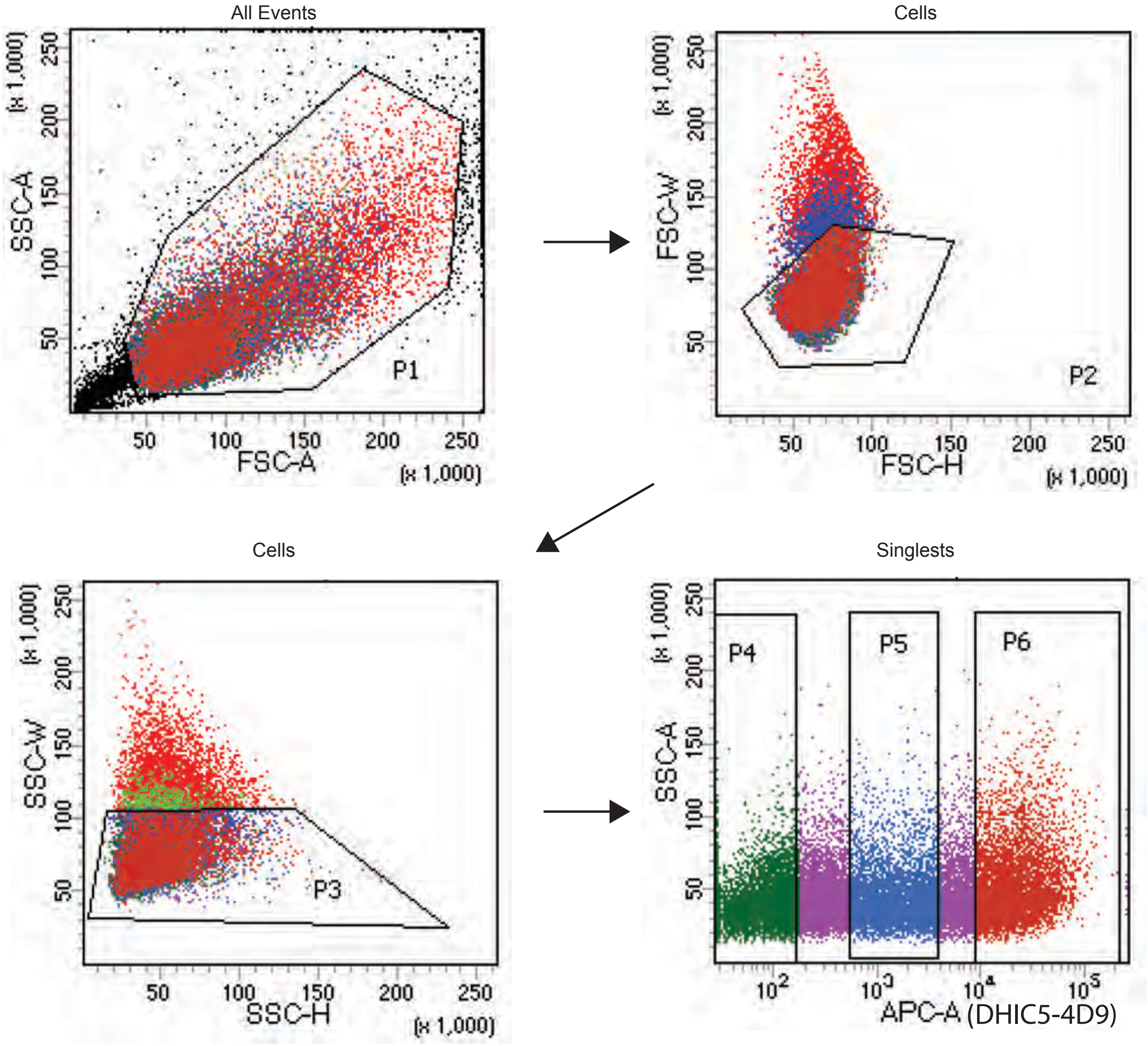

| Population | #Events | %Parent | %Total |
|---|---|---|---|
| All Events | 40,555 | #### | 100.0 |
| P1 | 30,000 | 74.0 | 74.0 |
| P2 | 24,812 | 82.7 | 61.2 |
| P3 | 25,436 | 84.8 | 62.7 |
| P2 AND P3 | 24,327 | 81.1 | 60.0 |
| P4 | 6,733 | 27.7 | 16.6 |
| P5 | 4,793 | 19.7 | 11.8 |
| P6 | 6,122 | 25.2 | 15.1 |

**Supplementary Figure 3**

Cell gating strategy for flow cytometry analysis and cell sorting experiments. APC-A was for DHIC5-4D9. Day49 H9 derived cholangiocytes were sorted for negative (P4) and positive (P6) populations (n=3).

a

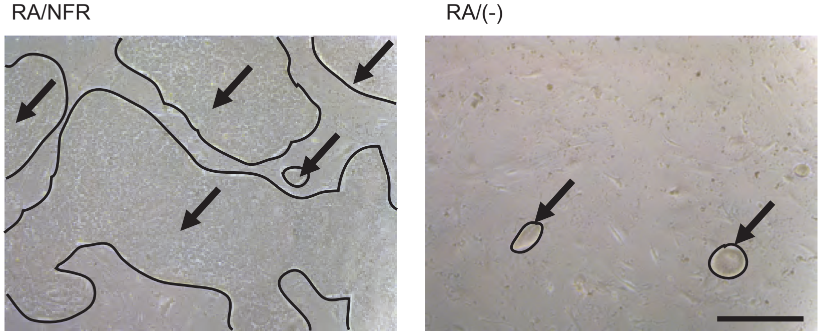

b

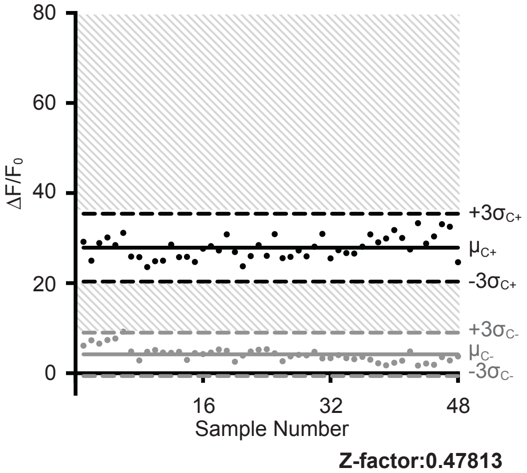

**Supplementary Figure 4**
**Characterization of hPSC-derived cholangiocytes.**

(a) Microphotographic images show the bright field images of day 49 cholangiocytes in the presence of indicated treatments. Line traces and arrows indicate the cholangiocyte colonies. Scale bar represents 500μm.

(b) The Z factors show the quality of the FLIPR assay measuring Apical Chloride Conductance (ACC) in 96 well plate with hPSC-derived cholangiocytes at day37.

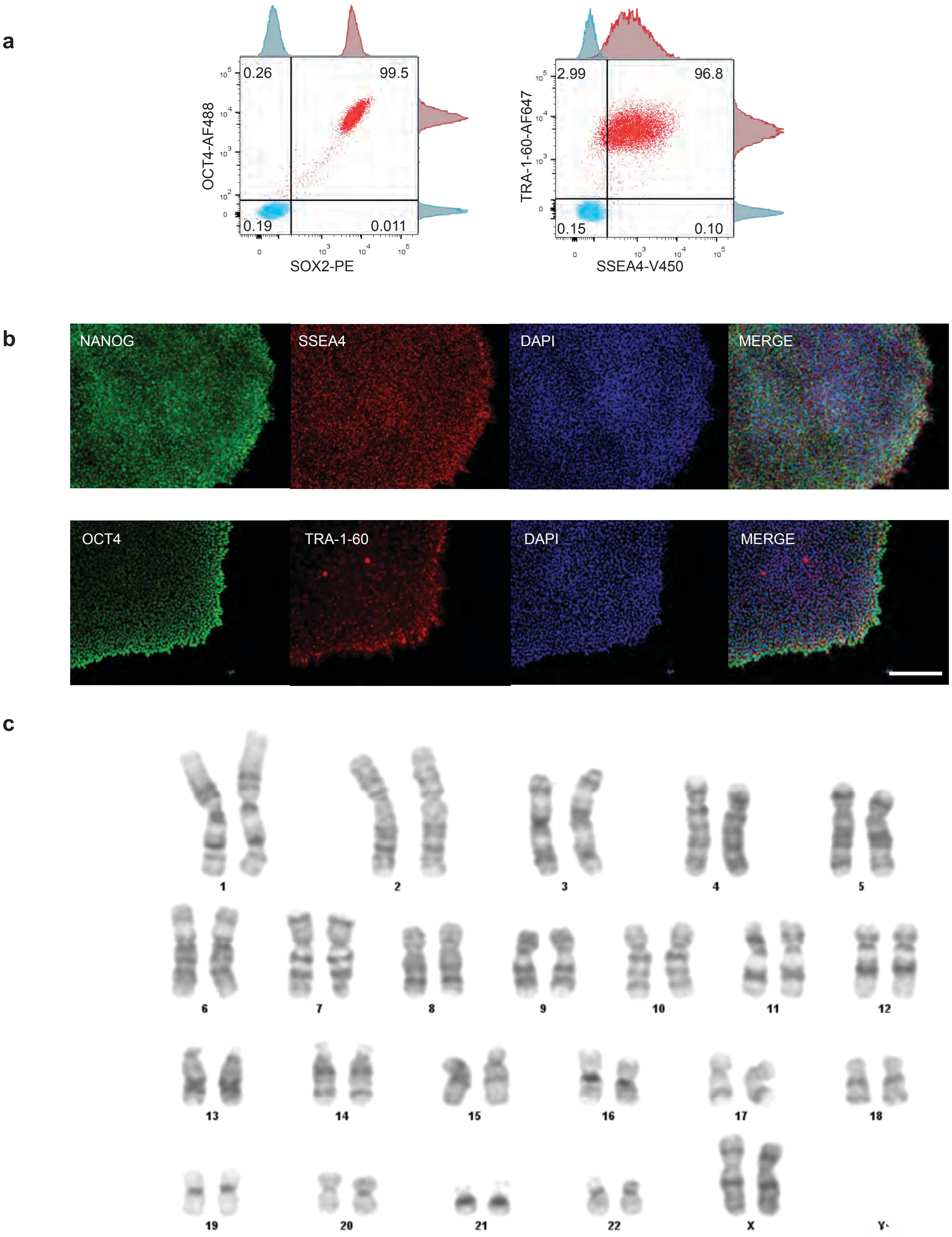
a
0.26
99.5
0.19
0.011
OCT4-AF488
SOX2-PE
2.99
96.8
0.15
0.10
TRA-1-60-AF647
SSEA4-V450
b
NANOG
SSEA4
DAPI
MERGE
OCT4
TRA-1-60
DAPI
MERGE
c
X
Y

**Supplementary Figure 5**
**Characterization of CCRM 45 (wild type) iPSCs.**

(a) Representative flow cytometric analysis showing OCT4, SOX2, TRA-1-60, and SSEA4 positivity.
(b) Immunostaining images of iPS colony demonstrating pluripotent markers NANOG, and SSEA4 in upper panel, OCT4 and TRA-1-60 in the lower panel. Scale bar represents 400μm.
(c) G-banding analysis shows normal karyotype.

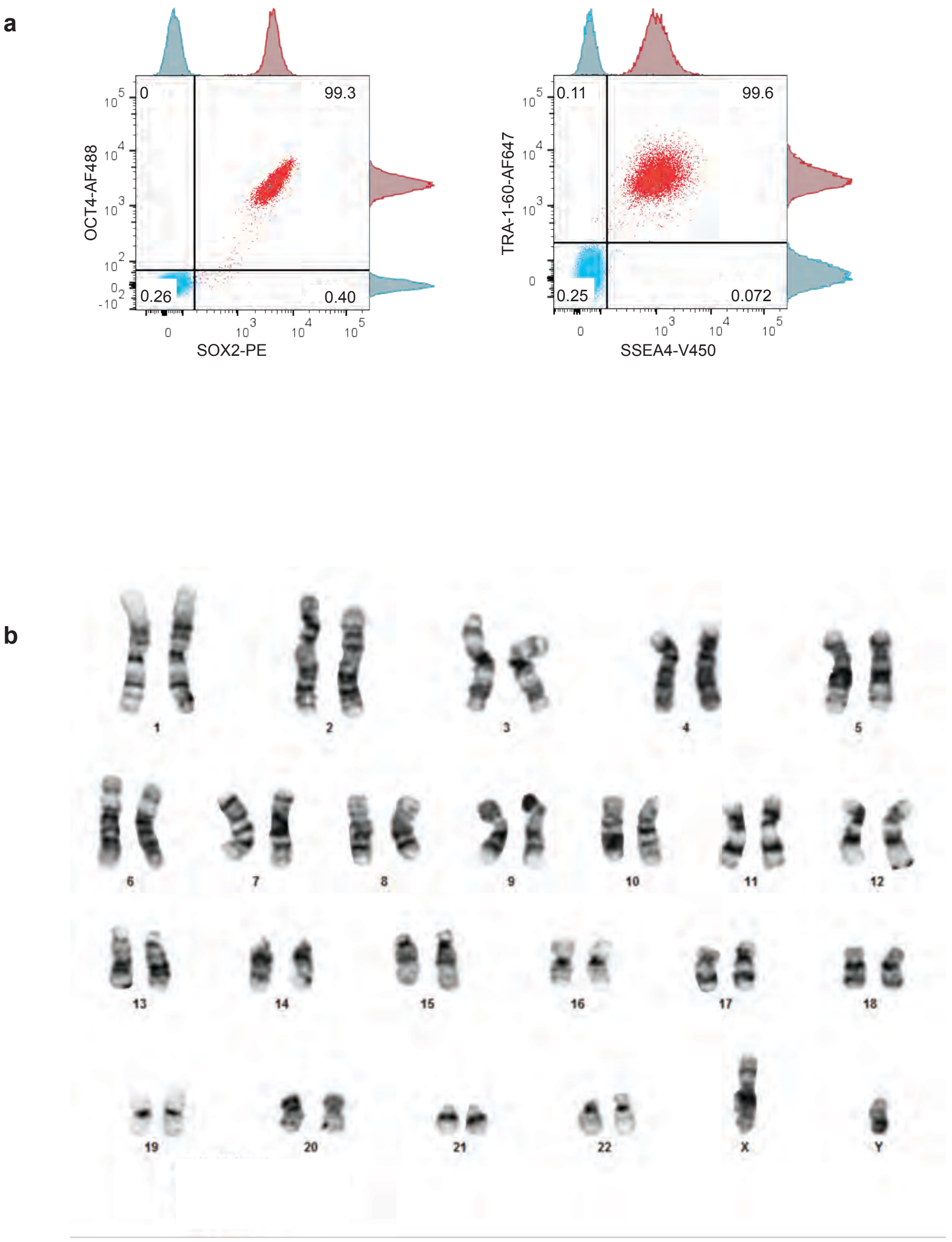

**Supplementary Figure 6**
**Characterization of CF9 (F508del) iPSCs.**

(a) Representative flow cytometric analysis showing OCT4, SOX2, TRA-1-60, and SSEA4 positivity.
(b) G-banding analysis shows normal karyotype.

**a**

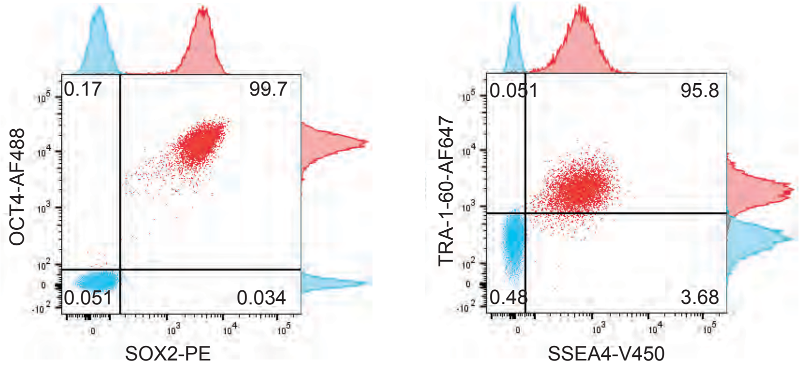
0.17
99.7
0.051
0.034
OCT4-AF488
SOX2-PE
0.051
95.8
0.48
3.68
TRA-1-60-AF647
SSEA4-V450

**b**

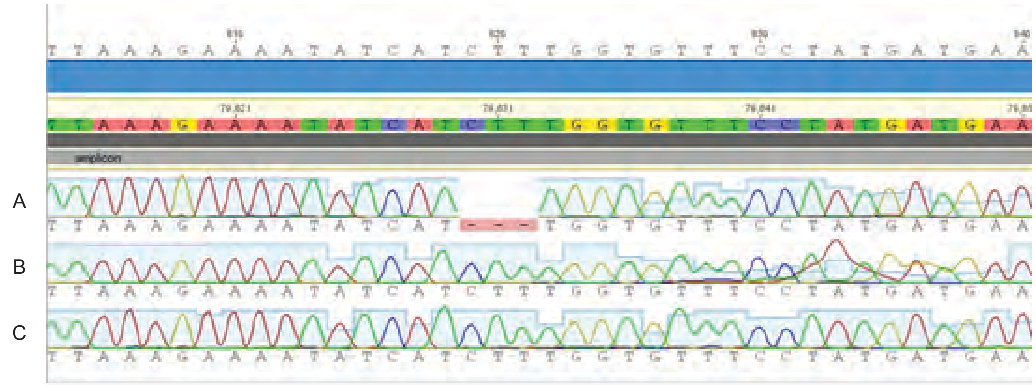
amplicon
A
B
C

**c**

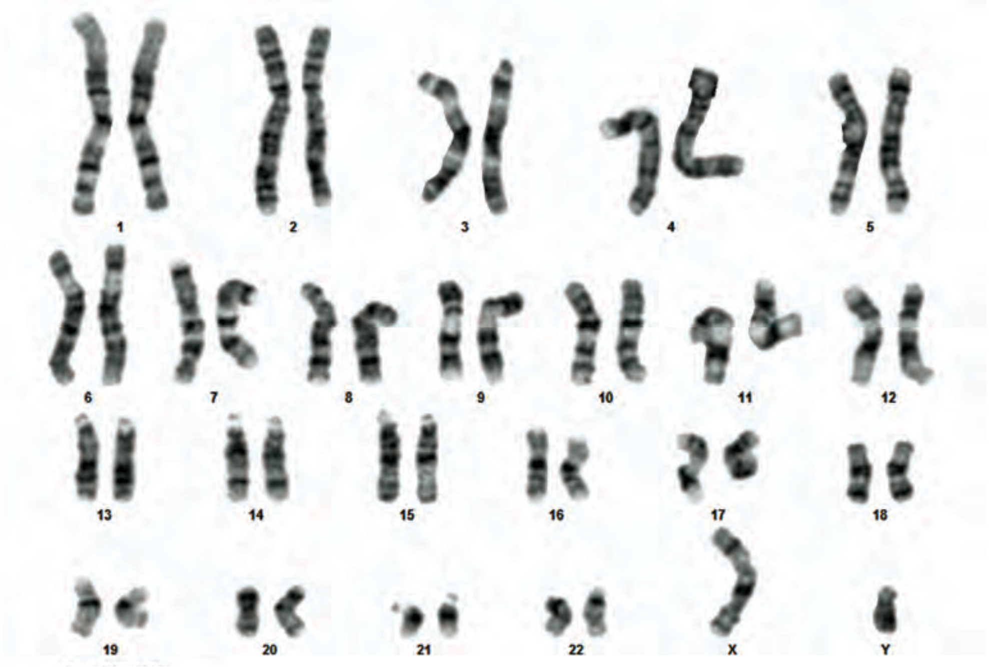
X
Y

**Supplementary Figure 7**
**Characterization of CF9GC (F508del mutation corrected) iPSCs.**

(a) Representative flow cytometric analysis showing OCT4, SOX2, TRA-1-60, and SSEA4 positivity.
(b) Sequencing of CFTR region of interest from (A) Genomic DNA extracted from CF9 showing cells homozygous for F508del. (B) Genomic DNA extracted from CF9GC corrected line showing cells homozygous for correction to wildtype. (C) Sequencing of (B) after TOPO cloning the CFTR PCR product showing the complete corrected sequence.
(c) G-banding analysis shows normal karyotype.

a

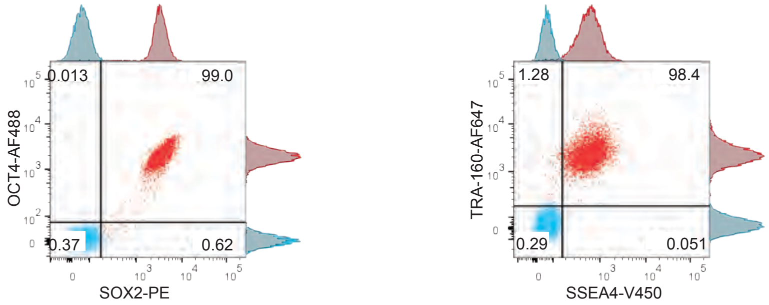

b

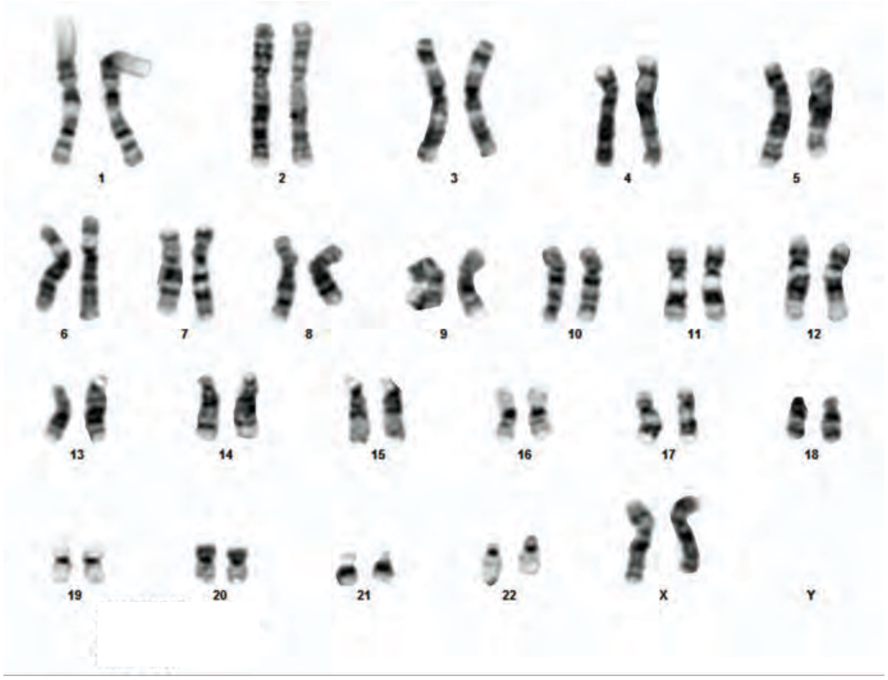

**Supplementary Figure 8**
**Characterization of CF10 (F508del) iPSCs.**

(a) Representative flow cytometric analysis showing OCT4, SOX2, TRA-1-60, and SSEA4 positivity.
(b) G-banding analysis shows normal karyotype.

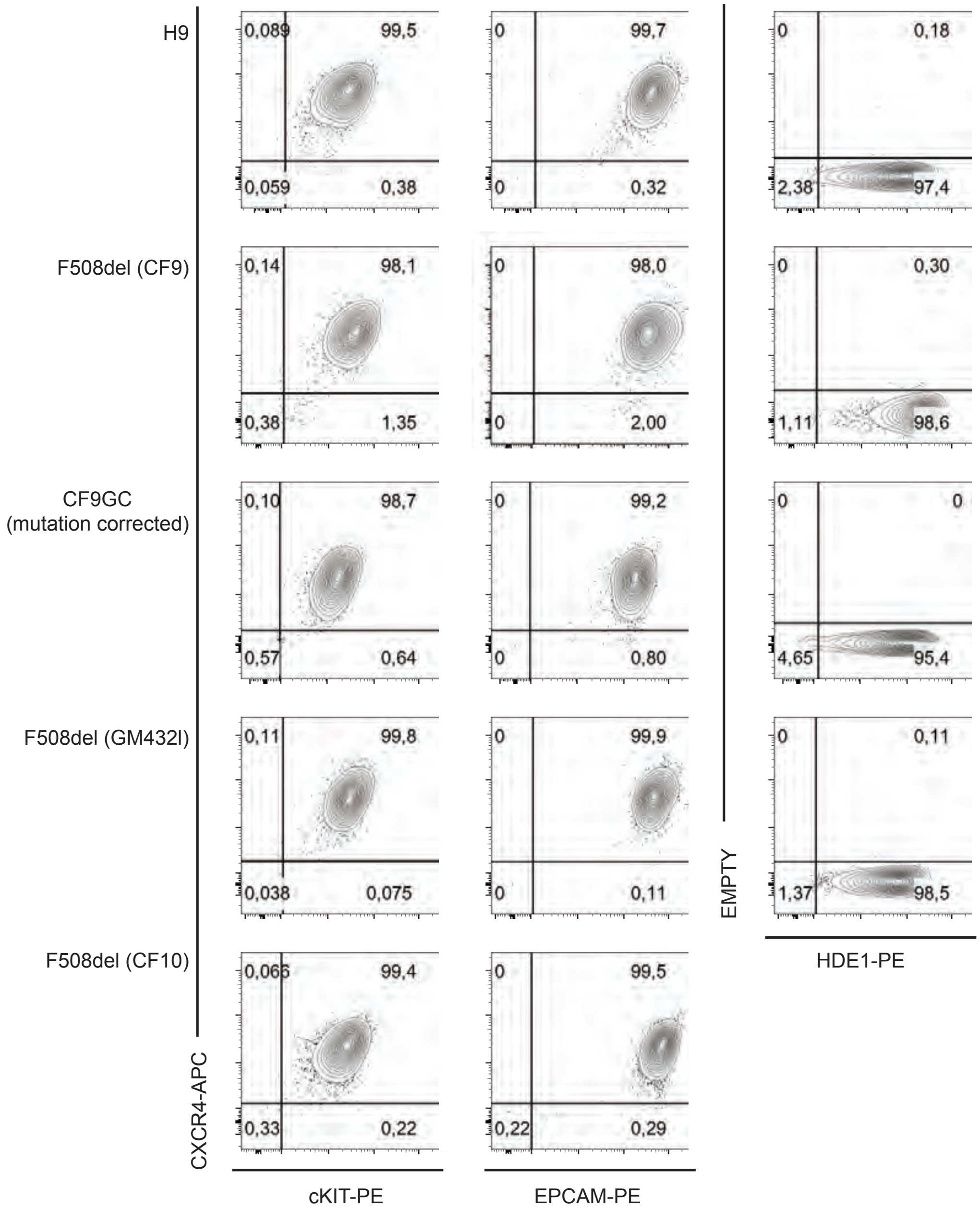

**Supplementary Figure 9**
**Characterization of hPSC-derived Endoderm.**
Representative flow cytometry analysis of day9 hPSC-derived endoderm from different cell lines.

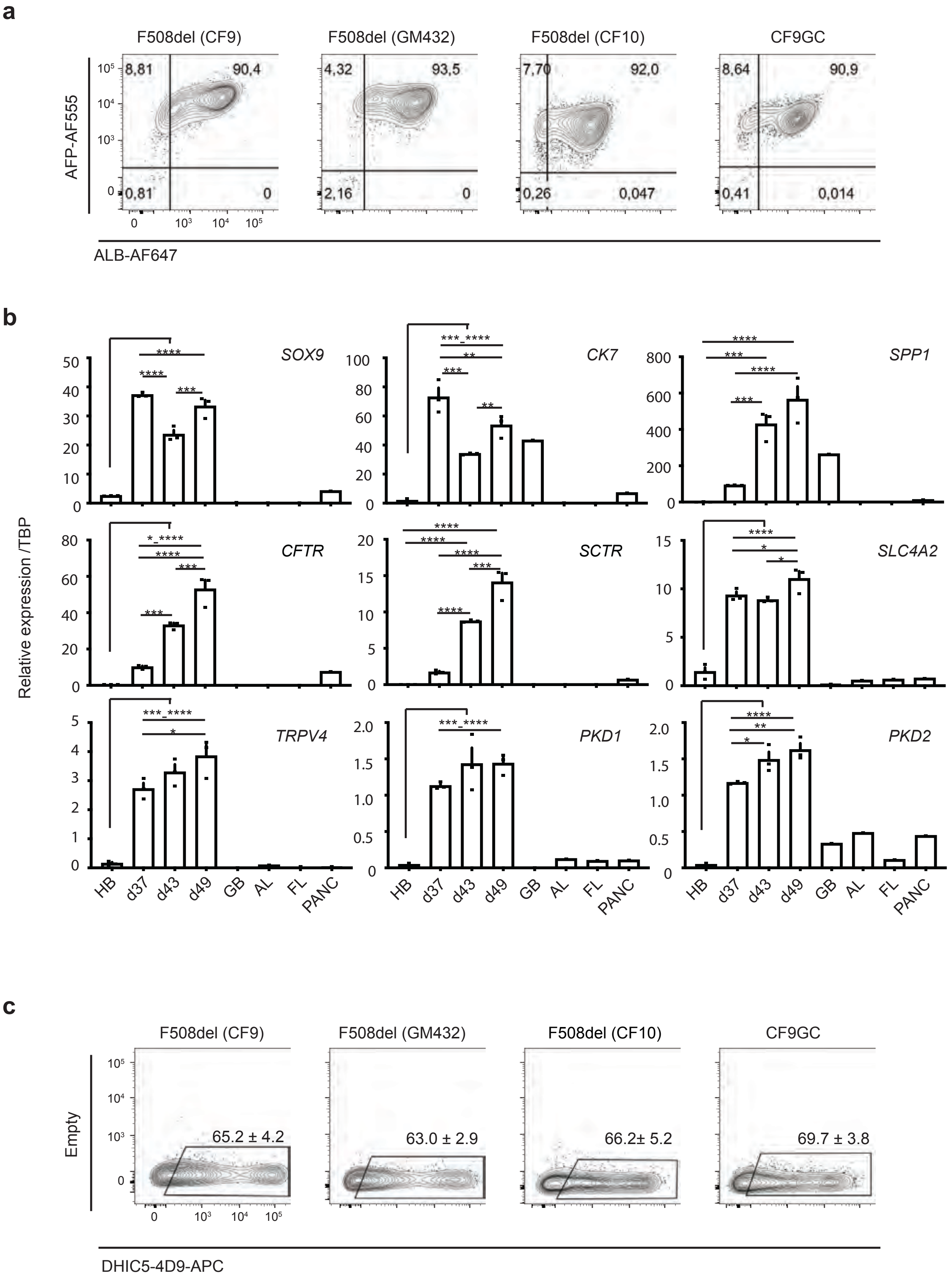
a
F508del (CF9)
F508del (GM432)
F508del (CF10)
CF9GC
AFP-AF555
ALB-AF647
8,81
90,4
0,81
0
4,32
93,5
2,16
0
7,70
92,0
0,26
0,047
8,64
90,9
0,41
0,014
b
SOX9
CK7
SPP1
CFTR
SCTR
SLC4A2
TRPV4
PKD1
PKD2
Relative expression /TBP
HB
d37
d43
d49
GB
AL
FL
PANC
c
F508del (CF9)
F508del (GM432)
F508del (CF10)
CF9GC
Empty
65.2 ± 4.2
63.0 ± 2.9
66.2± 5.2
69.7 ± 3.8
DHIC5-4D9-APC

**Supplementary Figure 10**
**Characterization of CF iPSC-derived cholangiocytes.**

(a) Representative flow cytometry analysis showing ALB and AFP positivity in day27-33 hPSC-derived hepatoblasts.
(b) qPCR analysis shows the expression of indicated genes in different stages of the CF9 (F508del) cholangiocyte differentiation. HB: hepatoblasts, GB: gall bladder, AL: adult liver, FL: fetal liver (n=3). Data are represented as mean ± SEM. $^{*}p \leq 0.05$, $^{**}p \leq 0.01$, $^{***}p \leq 0.001$, $^{****}p \leq 0.001$ one-way ANOVA.
(c) Representative flow cytometry analysis showing DHIC5-4D9 positivity in day49 hPSC derived cholangiocytes from day49-55 CF iPSC derived cholangiocytes (n=3-5). Data are represented as mean ± SEM.

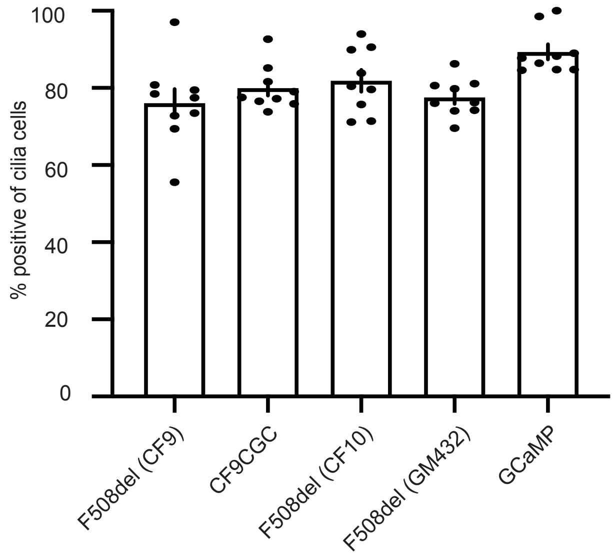

**Supplementary Figure 11**
**Cilia is highly positive among all cell lines.**
Quantification of cilia positive cells in day49 from different hPSC derived cholangiocytes (n=3). Data are represented as mean ± SEM.

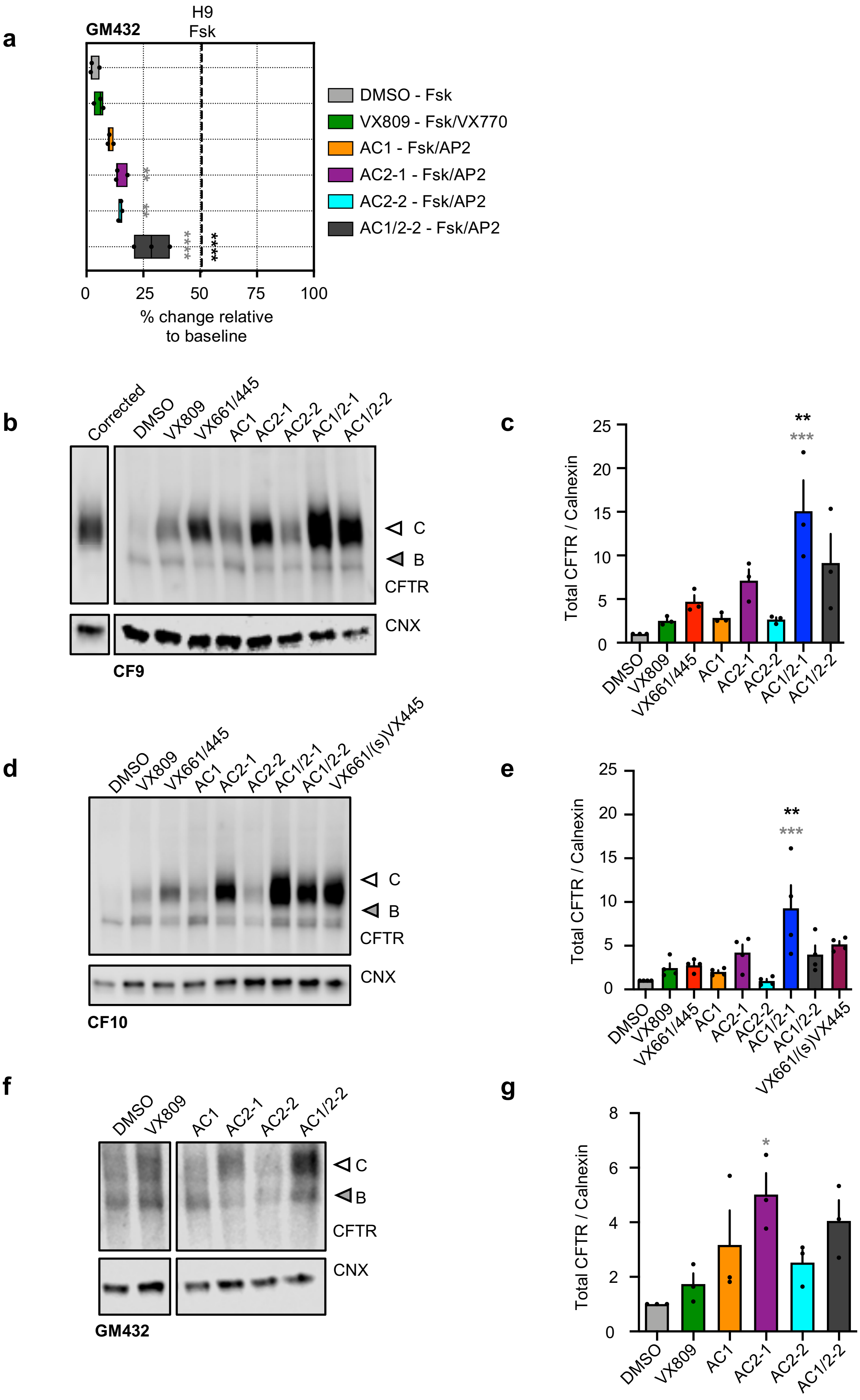
a
GM432
H9
Fsk
DMSO - Fsk
VX809 - Fsk/VX770
AC1 - Fsk/AP2
AC2-1 - Fsk/AP2
AC2-2 - Fsk/AP2
AC1/2-2 - Fsk/AP2
0
25
50
75
100
% change relative to baseline
b
Corrected
DMSO
VX809
VX661/445
AC1
AC2-1
AC2-2
AC1/2-1
AC1/2-2
C
B
CFTR
CNX
CF9
c
Total CFTR / Calnexin
d
DMSO
VX809
VX661/445
AC1
AC2-1
AC2-2
AC1/2-1
AC1/2-2
VX661/(s)VX445
C
B
CFTR
CNX
CF10
e
f
DMSO
VX809
AC1
AC2-1
AC2-2
AC1/2-2
C
B
CFTR
CNX
GM432
g
Total CFTR / Calnexin

**Supplementary Figure 12**
**Drug response in CF iPSC-derived cholangiocyte.**

(a) Representative % value of CFTR channel activity with an exposure of DMSO and CFTR modulators in GM432 iPSC-derived cholangiocytes normalized to DMSO. Dashed line indicates FSK response in H9-derived cholangiocytes. (n=3).
(b) Western blotting shows the immature (B) and mature (C) glycosylated CFTR bands after treatment with CFTR modulators in day49 F508del (CF9) cholangiocyte (right) and CF9GC cholangiocyte (left).
(c) Quantification of western blotting of the ratio of mature glycosylated CFTR protein with different CFTR modulators in day49 CF9 patient's iPSC-derived cholangiocytes (n=3).
(d) Western blotting shows the immature and mature glycosylated CFTR bands after treatment with CFTR modulators in day49 F508del (CF10) cholangiocyte.
(e) Quantification of western blotting of the ratio of mature glycosylated CFTR protein with different CFTR modulators in day49 CF9 patient's iPSC-derived cholangiocytes (n=4).
(f) Western blotting shows the immature and mature glycosylated CFTR bands after treatment with CFTR modulators in day49 F508del (GM432) cholangiocyte.
(g) Quantification of western blotting of the ratio of mature glycosylated CFTR protein with different CFTR modulators in day49 CF9 patient's iPSC-derived cholangiocytes (n=3). Data are represented as mean ± SEM. Grey and black stars show statistical significance to DMSO and VX809/VX770 respectively, one-way ANOVA. $^{*}p \leq 0.05$, $^{**}p \leq 0.01$, $^{***}p \leq 0.001$, $^{****}p \leq 0.0001$

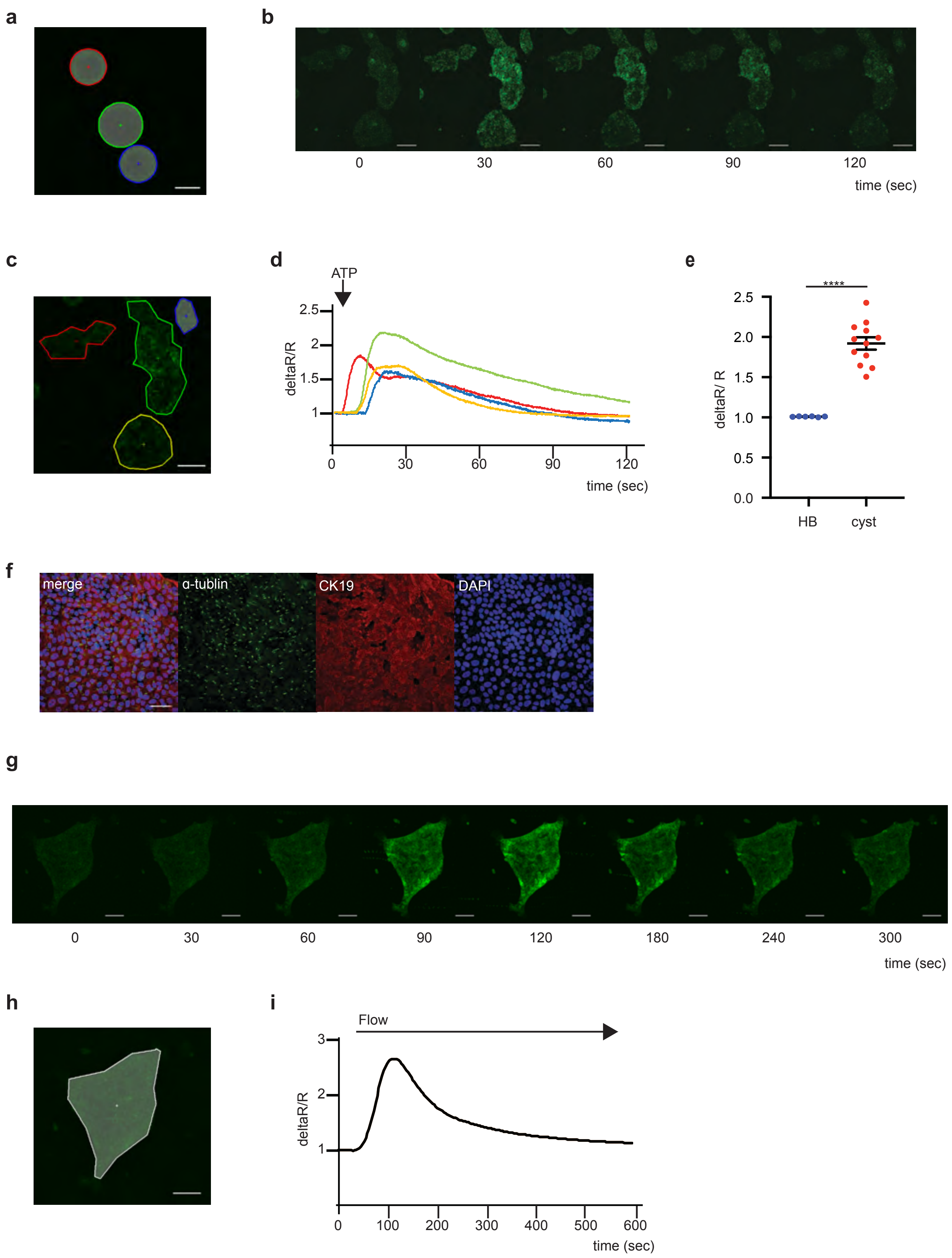
a
b
0
30
60
90
120
time (sec)
c
d
ATP
deltaR/R
2.5
2
1.5
1
0
30
60
90
120
time (sec)
e
****
deltaR/ R
2.5
2.0
1.5
1.0
0.5
0.0
HB
cyst
f
merge
α-tublin
CK19
DAPI
g
0
30
60
90
120
180
240
300
time (sec)
h
i
Flow
deltaR/R
3
2
1
0
100
200
300
400
500
600
time (sec)

**Supplementary Figure 13**
**Intracellular Calcium release in ciliated hPSCs derived cholangiocyte.**

(a) The region of interest (ROI)s correspond to GCaMP derived cholangiocyte cysts (3D) in main Figure 6(a). Scale bar represents 100 µm.
(b) Representative time lapse images of calcium influx in plated down 3D cholangiocyte cysts from GCaMP hESC in response to ATP. Scale bar represents 200µm.
(c) The ROIs mark the GCaMP derived plated down cholangiocyte cysts. Scale bar represents 100 µm.
(d) Representative traces showing the intracellular calcium release in plated down cholangiocyte cysts from GCaMP hESC in the response to ATP. The colors of the traces correspond to those for the ROIs in (c).
(e) Quantification of maximum fluorescent intensity representing intracellular calcium release in plated down 3D hepatoblast and 3D cholangiocyte cysts from GCaMP hESC in the response to ATP (HB n=4, chol n=3). Data are represented as mean ± SEM. $^{****}p \leq 0.0001$ two-tailed Student's t-test.
(f) Representative microscopic images of plate down 3D cholangiocyte cysts (GCaMP hESC) exhibiting primary cilia (green) and CK19 (red). Scale bar represents 50µm.
(g) Representative time lapse images of calcium influx in plated down 3D cholangiocyte cysts from GCaMP hESC in the response to flow. Scale bar represents 100µm.
(h) The ROIs mark the GCaMP derived plated down cholangiocyte cysts. Scale bar represents 100 µm.
(i) Representative trace of the intra cellular calcium release in plated down 3D cholangiocyte cysts from GCaMP hESC in the response to flow correspond to the ROI in (h).

### Supplementary Tables

**Supplementary Table 1.** Primary antibodies used for immunohistochemistry.

| Antibody | Company | Product Codes | IgG Species | Conjugate | Dilution |
|---|---|---|---|---|---|
| CFTR (24-1) | R and D | MAB25031 | Mouse | none | 1:200 |
| CFTR (13-1) | R and D | MAB1660 | Mouse | none | 1:200 |
| CK7 | Abcam | ab68459 | Rabbit | none | 1:200 |
| Acetylated α tubulin | Sigma-Aldrich | T7451 | Mouse | none | 1:800 |
| ZO-1 | Themo Fisher | 40-2200 | Rabbit | none | 1:400 |
| ARL13b | Proteintech | 17711 1-AP | Rabbit | none | 1:600 |
| CK19 | Abcam | ab52625 | Rabbit | none | 1:400 |
| Mitochondria | Millipore | MAB1273 | Mouse | none | 1:100 |
| AFP | DAKO | A0008 | Mouse | none | 1:2000 |
| SOX9 | Abcam | ab76997 | Mouse | none | 1:400 |
| ASBT (C14) | Santa Cruz | sc27493 | Goat | none | 1:50 |
| ALB | Bethyl | A80-129A | Goat | none | 1:200 |
| DHIC5-4D9 | Gift from oregon | | Mouse (IgM) | none | 1:20 |
| CD117 (c-KIT) | BD pharmingen | BD340529 | Mouse (IgG1) | PE | 1:50 |
| CD184 (CXCR4) | BD pharmingen | BD555976 | Mouse (IgG1) | APC | 1:50 |
| CD326 (EPCAM) | eBioscience | 12-9326-73 | Mouse (IgG1) | PE | 1:200 |
| SSEA4 | BD Horizon | 561156 | Mouse (IgG3) | V450 | 1:100 |
| TRA-1-60 | Biolegend | 330605 | Mouse (IgM) | Alexa Fluor 647 | 1:100 |
| OCT3/4 | BD pharmingen | 560791 | Mouse (IgG1) | Alexa Fluor 488 | 1:100 |
| SOX2 | BD pharmingen | 561556 | Mouse (IgG1) | PE | 1:100 |
| NANOG | BD pharmingen | 561506 | Mouse (IgG1) | PerCP-Cy5.5 | 1:100 |

**Supplementary Table 2.** Secondary antibodies used for immunohistochemistry.

| Antibody | Company | Product Codes | Dilution |
|---|---|---|---|
| IgG Donkey anti-Mouse Alexa488 | Invitrogen | A21202 | 1:400 |
| IgG Donkey anti-Rabbit Alexa555 | Invitrogen | A31572 | 1:400 |
| IgG Donkey anti-Mouse Alexa555 | Invitrogen | A31570 | 1:400 |
| IgG Donkey anti-Rabbit Alexa488 | Invitrogen | A21206 | 1:400 |
| IgG Donkey anti-Goat Alexa488 | Invitrogen | A11055 | 1:400 |
| IgM Goat anti-Mouse APC | Jackson ImmunoResearch | 115-136-075 | 1:200 |

**Supplementary Table 3.** Primers used for RT-PCR analysis.

| Gene | Sequences (Forward) | Sequences (Reverse) |
|---|---|---|
| **CFTR** | 5'-AGGACTATGGACACTTCGTGCCTT-3' | 5'-ATTTGGAACCAGCGCAGTGTTGAC-3' |
| **CK7** | 5'-AAGGATGCTCGTGCCAAG-3' | 5'-AGCTTCACGCTCATGAGTTC-3' |
| **SPP1** | 5'-CGAGGAGTTGAATGGTGCATA-3' | 5'-TCCAGCTGACTCGTTTCATAAC-3' |
| **TRPV4** | 5'-AGGTGAACTGGTCTCACTGG-3' | 5'-GCGAGAAGCCATAATACTGGTAG-3' |
| **PKD1** | 5'-GGACAAGGTGTGAGCCTGAG-3' | 5'-AGCTGGTAGACGTCCTCTGT-3' |
| **PKD2** | 5'-TTCCCAGATCAGTCATGGTTTAG-3' | 5'-CCTTCCATGCCTTCTGTAGATT-3' |
| **ALB** | 5'-GTGAAACACAAGCCCAAGGCAACA-3' | 5'-TCAGCCTTGCAGCACTTCTCTACA-3' |
| **AFP** | 5'-ACAGAGGAACAACTTGAGGCTGTC-3' | 5'-AGCAAAGCAGACTTCCTGTTCCTG-3' |
| **TMEM16A** | 5'-AAGTACTCGACGCTCCCGGCC-3' | 5'-ATAAGGAGTTCAGCAGCGTGCCC-3' |
| **AQP1** | 5'-TCTTCCGTGCCCTCATGTA-3' | 5'-CAAGCGAGTTCCCAGTCAG-3' |
| **SLC5A1** | 5'-TCAGGAGAGCCTATGACCTATT-3' | 5'-GGTGTCCGTCATCTTCATCTT-3' |
| **SLC4A2/AE2** | 5'-GGCATCTGTGCCCTCTTT-3' | 5'-TCCTGAATCTTGGGCTTGTC-3' |
| **ITPR3** | 5'-CGAGATGCTGCCCTTTGA-3' | 5'-CAGAGACGGGCAAACTTGA-3' |
| **P2YR** | 5'-GACTTCTTGTACGTGCTGACT-3' | 5'-GCTGCCATAGAGGTTCACAT-3' |
| **SOX9** | 5'-TGCATTTCCTCCTGCCTTTGCTTG-3' | 5'-GGGCACTTATTGGCTGCTGAAACA-3' |
| **SCTR** | 5'-TGCATCATGGCCAACTACTC-3' | 5'-AATCCCTGGAGGTACTTTCTTTC-3' |
| **TBP** | 5-'TGAGTTGCTCATACCGTGCTGCTA-3' | 5'-CCCTCAAACCAACTTGTCAACAGC-3' |

**Supplementary Table 4.** RNA control samples used for RT-PCR analysis.

| RNA | Source | Sex | Lot | Company | Product number |
|---|---|---|---|---|---|
| **human gall bladder** | **normal gall bladder from 34-years old** | **Female** | **A509245** | **BioChain** | **R1234118-10** |
| **human adult liver** | **normal livers pooled from 3 Asians (22-64-years old)** | **Male** | **1402003** | **Clonetech** | **636531** |
| **human fetal liver (Fig1b)** | **pooled from 63 spontaneously aborted fetus, aged 22-40 weeks** | **Male and Female** | **7030173** | **Clonetech** | **636540** |
| **human fetal liver (Fig2d,3e,Supp9b)** | **fetal liver from 24-weeks gestation** | **Male** | **1394** | **cell applications** | **1F21-50** |
| **human pancreas** | **normal pancreas from a 35-years old Caucasian** | **Male** | **1703157A** | **Clonetech** | **636577** |

### Description of Additional Supplementary Files

File Name: **Supplementary Video 1**
**Primary cilia expression in H9 monolayer cholangiocytes.**

File Name: **Supplementary Video 2**
**FSK induced swelling of H9-derived cysts.**

File Name: **Supplementary Video 3**
**ATP induced calcium signaling in GCaMP-derived 3D cholangiocyte cysts.**

File Name: **Supplementary Video 4**
**ATP induced calcium signaling in GCaMP-derived plated down cholangiocyte cysts.**

File Name: **Supplementary Video 5**
**Flow induced calcium signaling in GCaMP-derived cholangiocytes.**
